# Induction and *in silico* staging of human gastruloids with neural tube, segmented somites & advanced cell types

**DOI:** 10.1101/2024.02.10.579769

**Authors:** Nobuhiko Hamazaki, Wei Yang, Connor Kubo, Chengxiang Qiu, Beth K. Martin, Riddhiman K. Garge, Samuel G. Regalado, Eva Nichols, Choli Lee, Riza M. Daza, Sanjay Srivatsan, Jay Shendure

## Abstract

Embryonic organoids are emerging as powerful models for studying early mammalian development. For example, stem cell-derived ‘gastruloids’ form elongating structures containing all three germ layers^1–4^. However, although elongated, human gastruloids do not morphologically resemble post-implantation embryos. Here we show that a specific, discontinuous regimen of retinoic acid (RA) robustly induces human gastruloids with embryo-like morphological structures, including a neural tube and segmented somites. Single cell RNA-seq (sc-RNA-seq) further reveals that these human ‘RA-gastruloids’ contain more advanced cell types than conventional gastruloids, including neural crest cells, renal progenitor cells, skeletal muscle cells, and, rarely, neural progenitor cells. We apply a new approach to computationally stage human RA-gastruloids relative to somite-resolved mouse embryos, early human embryos and other gastruloid models, and find that the developmental stage of human RA-gastruloids is comparable to that of E9.5 mouse embryos, although some cell types show greater or lesser progression. We chemically perturb WNT and BMP signaling in human RA-gastruloids and find that these signaling pathways regulate somite patterning and neural tube length, respectively, while genetic perturbation of the transcription factors PAX3 and TBX6 markedly compromises the formation of neural crest and somites/renal cells, respectively. Human RA-gastruloids complement other embryonic organoids in serving as a simple, robust and screenable model for decoding early human embryogenesis.

## INTRODUCTION

The molecular and cellular biology of early human development is of fundamental interest. However, due to ethical, technical and practical difficulties, opportunities to study *in vivo* post-implantation early human development are limited. To address this in part, several systems for *ex vivo* culture of embryos have been established, *e.g.* extending their development by providing an attachment substrate or extracellular matrix such as 3D Matrigel^5–7^. However, these approaches still require fertilized human embryos as a starting point.

A different approach to study early human development involves *in vitro* models that begin with pluripotent stem cells (PSCs)^8–13^. For example, ‘gastruloids’ leverage the self-organizing potential of PSCs^1–4,14^ and are distinguished by the formation of derivatives of all three germ layers, as well as by the specification of an anteroposterior axis^3^. Recently, the addition of Matrigel during the induction of mouse gastruloids resulted in morphological features even more characteristic of *in vivo* development, including a neural tube and segmented somites^2,15^, while treatment of PSC aggregates with a presomitic mesoderm-promoting cocktail was recently shown to result in human ‘somitoids’^16^. However, human gastruloids that robustly form both a neural tube and segmented somites have yet to be demonstrated. Furthermore, it remains unclear how to temporally ‘stage’ the progression of these complex *in vitro* models, in relation to one another as well as to *in vivo* mammalian development.

We investigated differences between mouse and human gastruloids generated with published protocols, which led us to hypothesize that neuromesodermal progenitor (NMPs) are biased towards somitic rather than neural fates in previously described human gastruloids, relative to both mouse gastruloids and *in vivo* development. In seeking to correct this, we discovered that a discontinuous regimen of retinoic acid (RA) and Matrigel is sufficient to induce embryo-like morphological features in human gastruloids. Specifically, these human ‘RA-gastruloids’ robustly form both a neural tube and segmented somite-like structures. Furthermore, sc-RNA-seq revealed that human RA-gastruloids include more advanced cell types than either human or mouse gastruloids, including neural crest cells, renal progenitor cells, skeletal muscle cells, and neural progenitor cells. We developed and applied a new method to computationally stage human RA-gastruloids and other mammalian gastruloid models in relation to *in vivo* human and mouse development. We also tested the effects of chemically perturbing various signaling pathways, or genetically perturbing key transcription factors, on the emergence of specific structures and cell types in human RA-gastruloids. Human RA-gastruloids form robustly and are highly screenable, further advancing the prospects of entirely *in vitro* models to recapitulate key aspects of early human development.

## RESULTS

### Comparative analysis of the dynamics of human vs. mouse gastruloid formation

We set out to induce human gastruloids with more advanced morphological features than published protocols^1^. Specifically, we sought to demonstrate the elongation of a neural tube flanked by segmented somites. Our early experiments simply attempted to adopt modifications that resulted in mouse gastruloids with trunk-like structures (TLS)^15^, *e.g.* the addition of Matrigel. However, unlike mouse TLS, the addition of Matrigel did not alter the morphology of human gastruloids, although it did substantially increase the success rate of elongation (**Fig. S1**; **Table S1**). We hypothesized that differences between human and mouse gastruloid composition might underlie their differential responses to Matrigel. Human gastruloids without Matrigel were previously characterized by Tomo-seq^17^, a method that profiles RNA on a series of cryosections, but not by sc-RNA-seq^1^. We therefore performed sc-RNA-seq on such human gastruloids^17^, *i.e.* without Matrigel, at 24, 48, 72 and 96 hrs after induction (**Fig. 1a-b**). After filtering low-quality cells and doublets, we obtained profiles for ∼44,000 cells across these four timepoints, which were subjected to dimensionality reduction, unsupervised Louvain clustering, and cell type annotation (**Fig. 1c**; **Table S2**).

**Figure 1.**
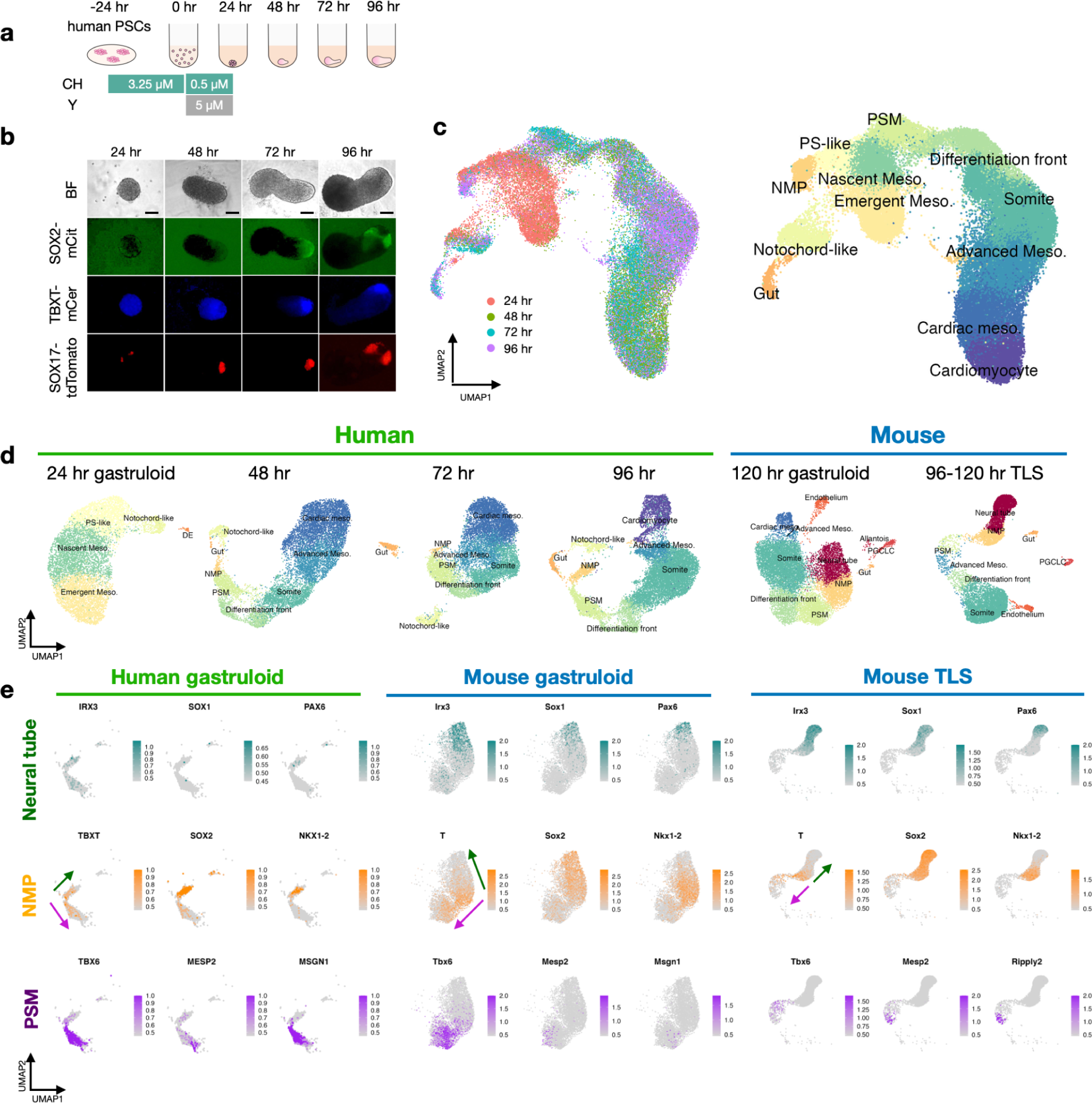
Time-resolved, single cell transcriptional profiling of human gastruloids. **a**, Schematic of human gastruloid protocol^1^. CH, CHIR99021; Y, Y-27632. **b**, Representative images of RUES2-GLR^26^ hPSC-derived human gastruloids. SOX2-mCit, pluripotent and ectoderm marker; TBXT-mCer, mesoderm marker; SOX17-tdTomato, endoderm marker. Scale bar, 100 µm. **c**, Integrated UMAP of ∼44,000 sc-RNA-seq profiles from four timepoints of human gastruloid development, colored by timepoint (left) or cell type annotation (right). NMP, neural mesodermal progenitor; PSM, presomitic mesoderm; PS, primitive streak. **d**, UMAP projection of sc-RNA-seq profiles from individual timepoints of human gastruloids (generated by this study based on published protocols^1^) or published data from mouse gastruloids^2^ or mouse TLS^15^. PGCLC, primordial germ cell-like cell; DE, definitive endoderm;. **e**, Normalized expression of marker genes for neural tube (upper row), NMPs (Middle row), or PSM (lower row) in UMAP projections of sc-RNA-seq profiles of neural tube, NMP, or PSM cells from human gastruloids at 96 hrs, mouse gastruloids at 120 hrs^2^, or mouse TLS at 120 hrs^15^. Arrows represent putative differentiation of NMPs towards neural tube (green) and PSM (purple) fates, respectively. The key point is that NMP-like and PSM-like cells are detected in all three models, but neural tube-like cells are detected only in the two mouse models.

We sought to follow the emergence of key cell populations across this time course of human gastruloid formation (**Fig. 1d**; **Fig. S2-S5**). At 24 hrs after seeding, the bulk of cells form a continuum in a Uniform Manifold Approximation and Projection (UMAP) representation, with heterogeneous expression of *TBXT*, *CDX2*, *TBX6*, *SOX2*, *CER1*, *FOXF1*, *HAND1*, *OSR1*, *GATA6* and other markers (**Fig. 1d**; **Fig. S2; Table S3**). Distinct populations of *FOXA2*+ cells were either *NOTO*+ or *SOX17*+, presumably corresponding to axial mesoderm and definitive endoderm equivalents, respectively (**Fig. S2**). For annotation of the bulk of cells, integration with these 24 hrs sc-RNA-seq data from a recently reported Carnegie Stage (CS7) human embryo^18^ was informative, as these mapped to a continuum of similarity with the *in vivo* human primitive streak, nascent mesoderm and emergent mesoderm (**Fig. S6a**). However, extraembryonic, hematopoietic endothelial, and primordial germ cell (PGC) equivalents were missing from the human gastruloid, as were epiblast and non-neural ectoderm equivalents.

Integrated projections of the 48 to 96 hrs human gastruloid and CS7 human embryo profiles were less informative, potentially because the gastruloids had advanced beyond this *in vivo* stage (**Fig. S6b-d**). However, through integration with mouse gastruloid and TLS data^15^ (**Fig. S7**), we were able to putatively annotate more differentiated subpopulations of cells at these later timepoints, including a continuum from NMPs (*TBXT*+; *SOX2*+; *NKX1-2*+) to presomitic mesoderm (*TBX6+*) to a differentiation front (*MESP2*+; *RIPPLY2*+; *RIPPLY1*+) giving rise to differentiated somites (*FST+*; *PAX3*+); and additionally, advanced (*OSR1+*; *WT1-)* and cardiac mesoderm (*HAND1*+; *NKX2-5*+; *TNNT2+*) (**Fig. 1d**; **Fig. S3-S5**). Of note, when we integrated human gastruloid profiles across all four timepoints, the subpopulations annotated as advanced and cardiac mesoderm at 48 to 96 hrs were not easily relatable to nascent and emerging mesoderm at 24 hrs (**Fig. 1c**), raising the possibility of rapid changes that would benefit from denser sampling across the 24 to 48 hrs interval. Axial mesoderm or notochord-like cells (*NOTO*+) were also detected at these later timepoints (**Fig. 1d**; **Fig. S3-S5**). Through integration with mouse E6.5-E8.5 data^19^, we were also able to track the gut/endoderm-like subpopulation through 96 hrs (**Fig. S8**), although *SOX17* expression was inconsistent in these cells (**Fig. S3-S5**).

We next sought to explore differences between human and mouse gastruloids. In embryos, the posterior neural tube arises from bipotential stem cells called neuromesodermal progenitors (NMPs)^20–24^. Although NMP-like cells were detected in mouse gastruloids^2^, mouse TLS^15^ and human gastruloids^1^, and indeed appear to be a source of presomitic mesoderm in all three cases, we observed the emergence of neural tube-like cells (*IRX3+*; *SOX1*+; *PAX6+*) only in mouse gastruloids and mouse TLS, and not in human gastruloids (**Fig. 1d-e**; **Table S2**). We therefore hypothesized that a bias in differentiation potential of NMPs towards mesodermal fates might underlie the failure of human gastruloids to generate an elongated neural tube, and, given that they might be induced by neural tube signaling^22,25^, perhaps segmented somites as well.

### A discontinuous regimen of retinoic acid robustly induces a neural tube and segmented somites in human gastruloids

To facilitate the differentiation of neural lineages from NMPs, we sought to supply additional signaling cues, presumably insufficiently provided in the human gastruloid system. Retinoic acid (RA) is one such signaling molecule that can induce neural cell fates from NMPs, both *in vivo* and *in vitro*^24,27–33^. Additionally, RA plays a crucial role in the patterning of somites as an anterior signal^34,35^. Thus, in addition to the 10% Matrigel (which, on its own, failed to induce a neural tube or segmented somites; **Fig. S1**; **Table S1**), we explored adding various concentrations of RA to the gastruloid induction medium (**Fig. S9a-b**). Encouragingly, SOX2-mCit intensities increased at 24 hrs in a dose-dependent manner (**Fig. S9c**). However, although gastruloid elongation was enhanced at RA concentrations ranging from 100 nM to 1µM, we did not observe neural tube formation nor somite segmentation.

We speculated that continuous RA exposure might perturb the differentiation of other cell types, *e.g.* the somitic and cardiac lineages, between 24 and 48 hrs. We, therefore, attempted a similar experiment, but we withdrew RA at 24 hrs and then added it back at 48 hrs together with 10% Matrigel (**Fig. S10a**). Remarkably, this discontinuous regimen induced the elongation of SOX2-mCit-positive neural tube-like structures as well as the formation of primitive segmentations of apparent somites (**Fig. S10b**). Building on this finding, we sought to optimize the number of cells used in the initial seeding of gastruloids (**Fig. S10c**). With this discontinuous regimen of RA and a larger number of initial cells, gastruloids began to exhibit multiple, segmented somites, together with a neural tube-like structure along an A-P axis (**Fig. 2a-b**). These structures formed robustly under these conditions, with 89% of elongated gastruloids exhibiting both segmented somite and neural tube-like structures across five independent experiments (**Fig. 2c**).

**Figure 2.**
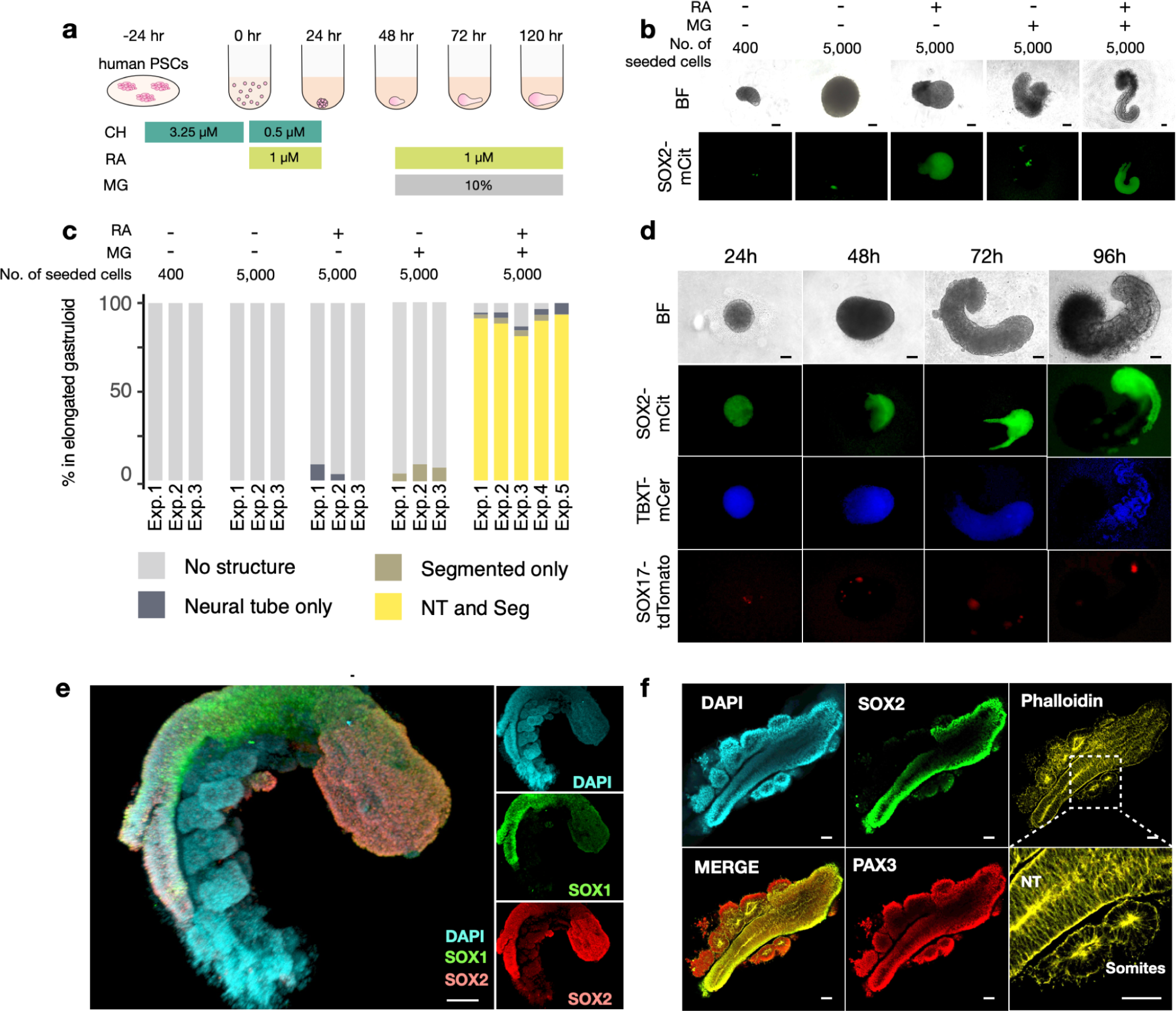
Robust induction of somites and neural tube in human gastruloids via a discontinuous regimen of retinoic acid. **a**, Schematic of human RA-gastruloid protocol. RA, retinoic acid; MG, Matrigel; CH; CHIR99021. RA is applied for the first 24 hrs after induction, then withdrawn, then added back at 48 hrs along with 10% Matrigel. **b**, Representative images of 120 hr human gastruloids with vs. without RA and MG, from 400 or 5,000 seeding cells. Scale bar, 100 µm. **c**, Quantification of frequency of neural tube elongation and somite segmentation. NT, Neural tube; Seg, segmented somites. **d**, Representative images of developing human RA-gastruloids. Scale bar, 100 µm. N=768 (96 x 8 plates) human RA-gastruloids showed similar morphology and expression patterns of marker genes. **e**, 3D projections of immunostained 120 hr RA-gastruloids. SOX1+, SOX2+ region corresponds to neural tube-like structure, flanked by somite-like structures. N=12 human RA-gastruloids showed similar morphology and expression patterns of marker genes. **f**, Confocal section of immunostained 120 hr RA-gastruloid. Phalloidin staining shows the apical accumulation of F-actin in SOX2+, PAX3+ neural tube and PAX3+ somites. Magnified region is indicated by a dotted square. Scale bar, 100 µm. N=9 human RA-gastruloids showed similar morphology and expression patterns of marker genes.

The spatial distribution of germ-layer specific markers in these RUES2-GLR^26^-derived gastruloids was consistent with the presence of these structures (**Fig. 2d**). Immunostaining confirmed that the neural tube-like structures were SOX2+/SOX1+ (**Fig. 2e-f**). We also immunostained these gastruloids with antibodies to PAX3, a marker of epithelialized somites^36^, and phalloidin, which stains apical actin^37^. The somite-like structures were PAX3+ and exhibited an accumulation of F-actin on their apical sides, as did the SOX2+ neural tube-like structures (**Fig. 2f**). This strongly suggests that these structures possess epithelial features, as such asymmetric accumulation is also seen in embryonic somites and neural tubes *in vivo*^38–40^. Together, these results show that a discontinuous regimen of RA can robustly induce embryo-like morphological structures, including a neural tube and segmented somites, in human gastruloids. We hereafter refer to these as ‘human RA-gastruloids’.

### Transcriptional profiling of human RA-gastruloids

To identify the cell types present, we applied sc-RNA-seq to human RA-gastruloids. Clustering and annotation of 5,347 and 18,324 single cell profiles identified 9 and 12 cell types at 96 and 120 hrs, respectively (**Fig. 3a**; **Fig. S11a**). Although a subset of the cell types observed in conventional human gastruloids were observed (cardiac mesoderm, differentiation front, somites, gut), we additionally identified cells resembling neural tube (*PAX6*+; *SOX1*+), neural crest (*FOXD3*+; *SOX10*+), neural progenitors (*ONECUT1*+; *ONECUT2*+), intermediate mesoderm (*WT1*+; *OSR1*+), and renal epithelium (*LHX1*+; *PAX2*+) at 96 hrs and, additionally, myocytes (*NEB*+), cardiomyocytes (*TNNT2*+) and endothelium (*PLVAP*+) at 120 hrs, in human RA-gastruloids (**Fig. 3a**; **Fig. S11b**).

**Figure 3.**
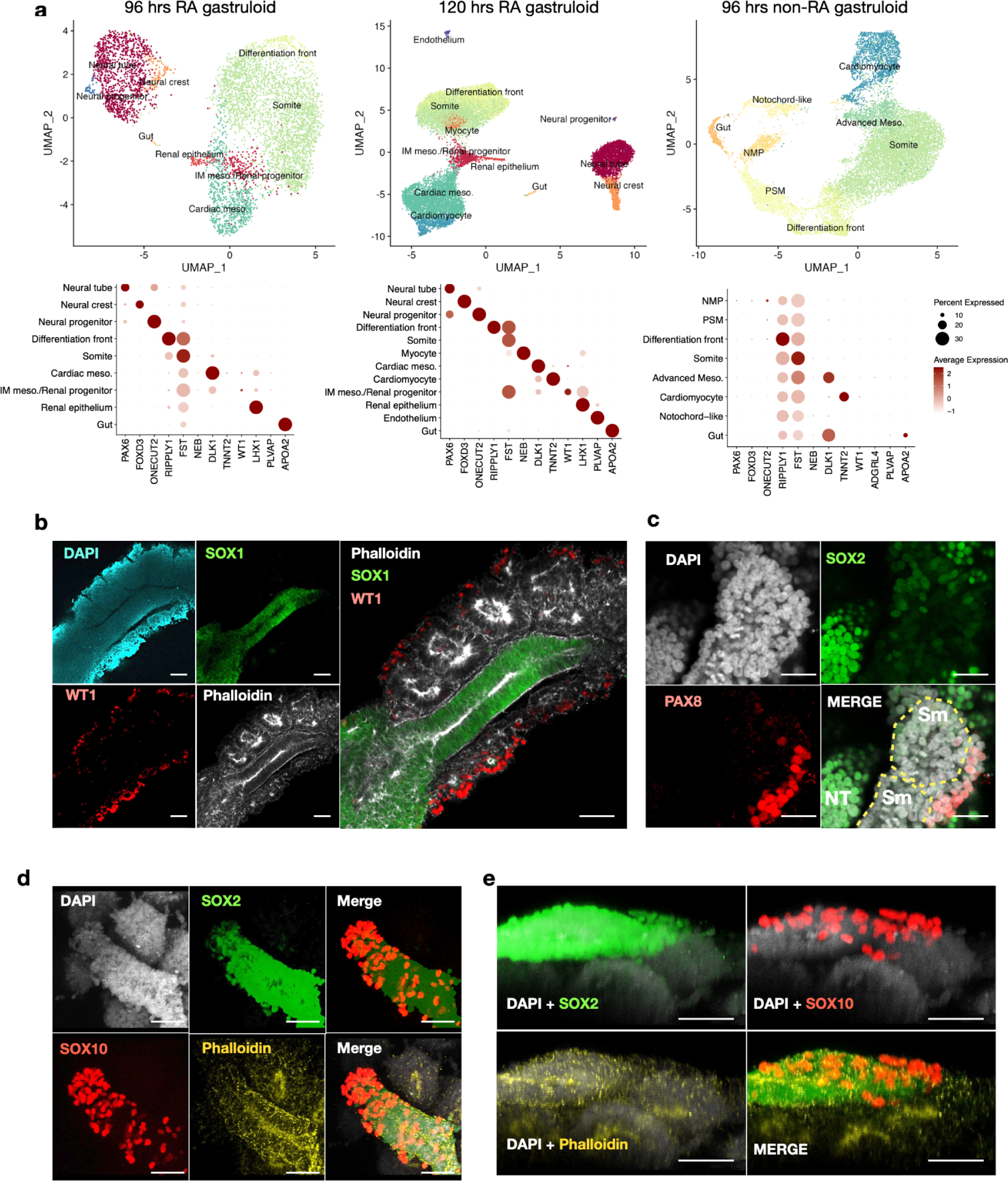
Induction of neural crest, intermediate mesoderm and other advanced cell types in human RA-gastruloids. **a**, (upper) UMAP of sc-RNA-seq profiles from 96 hr non-RA-gastruloids, 96 hr RA-gastruloids, and 120 hr RA-gastruloids, colored and labeled by cell type. PSM, presomitic mesoderm; DF, differentiation front; NMP, neuromesodermal progenitor; IM meso., intermediate mesoderm. (lower) Bubble plot of expression of marker genes for each annotated cell type. **b-c**, Immunostaining of intermediate mesoderm cells and renal epithelium in 120 hrs RA-gastruloids. **b**, Immunostaining of 120 hr RA-gastruloid with anti-WT1 antibody (red, intermediate mesoderm cells), anti-SOX1 antibody (green, neural tube), phalloidin (white, F-actin) and DAPI (cyan, nuclear). *WT1*+ intermediate mesoderm-like cells appear lateral to the phalloidin-stained somites. Scale bar, 100 µm. **c**, Immunostaining of 120 hr RA-gastruloid with anti-PAX8 antibody (red, renal epithelium), anti-SOX2 antibody (green, neural tube), and DAPI (cyan, nuclear). Scale bar, 100 µm. N=4 (c) and N=3 (d) showed similar morphology and expression pattern of marker genes. Sm, somite; NT, neural tube. **d-e**, Immunostaining of neural crest-like cells in 120 hrs RA-gastruloid. **d**, 3D projection of somite and neural tube in 120 hr RA-gastruloid stained with anti-SOX2 antibody (green, neural cells), anti-SOX10 antibody (red, neural crest), phalloidin (yellow, F-actin) and DAPI (white, nuclear). **e**, Reconstructed lateral views of RA-gastruloids shown in panel (**d**). *SOX10*+ neural crest-like cells occupy the putative dorsal surface of the *SOX2*+ neural tube-like structure. Scale bar, 50 µm. N=11 showed similar morphology and expression pattern of marker genes.

We were surprised to observe many of these additional cell types, *e.g.* intermediate mesoderm (IMM) (*OSR1*+, *WT1*+) and renal epithelium (RE) (*PAX8+, ADGRL4+*), because they are absent not only from the original human gastruloid model, but also from the mouse gastruloid and TLS models^1,2,15^. During *in vivo* development, IMM arises between the somites and lateral plate mesoderm along the mediolateral axis, analogous to their topological distribution in the UMAPs of human RA-gastruloids (**Fig. 3a**). To investigate the spatial distribution of the IMM-like cells, we immunostained human RA-gastruloids with anti-WT1 antibody or anti-PAX8. Consistent with expectation, the IMM-like *WT1*+ cells were located lateral to somite structures (**Fig. 3b**).

Neural crest cells (*SOX10*+, *FOXD3*+) were also unexpected, as these are similarly absent from all mammalian gastruloid models described to date. Neural crest cells are a multipotent cell type, arising at the dorsal aspect of the neural tube and migrating throughout the embryo, giving rise to numerous mesodermal and ectodermal derivatives^41^. To visualize the spatial distribution of neural crest cells in relation to the neural tube, we immunostained human RA-gastruloids with anti-SOX10 antibody (**Fig. 3c-d**). Remarkably, neural crest cells were observed on the surface of the neural tube-like structure (**Fig. 3c**). Moreover, the z-stack analysis showed that these neural crest cells were preferentially located on one side of the neural tube, potentially the dorsal equivalent, suggesting that the spatial patterning of neural crest cells may be recapitulated in the human RA-gastruloid system (**Fig. 3d**).

To investigate the potential spatial patterning of neuroectoderm more deeply, we reanalyzed neural tube, neural crest, and neural progenitor cells on their own, and then examined the expression patterns of marker genes in the resulting embedding (**Fig. S12a-d**; **Table S4**). A large proportion of neural tube cells expressed dorsal neural tube or roof plate markers, including *PAX3*. In contrast, ventral neural tube or floor plate markers are not expressed (although *SP8*, which is present at the boundary of the dorsal and ventral regions of the neural tube in normal development^42^, is detected). Together with the asymmetric appearance of neural crest cells on one surface of the RA-gastruloids’ neural tubes, these results suggest that the human RA-gastruloids are dorsally biased. We speculate that the incomplete establishment of the D-V axis is due to the lack of a Sonic hedgehog (SHH)-secreting notochord^43^.

Is any aspect of neurogenesis ongoing in human RA-gastruloids? Neural progenitor cells arise from a stem cell pool of radial glial cells in the neural tube. We analyzed expression patterns of HES genes, which are markers of radial glial cells^44^, in RA-gastruloids (**Fig. S12e**). Among cells annotated as neural tube, we observe a subpopulation of radial glial cell-like cells (*HES1*+*, HES3*-, *HES6*+), and, separately, a subpopulation of neural progenitor-like cells (*HES5*+, *POU3F1*+*, ONECUT1*+). To confirm this, we immunostained neural progenitor cells with anti-POU3F1 antibody, and found the neural progenitor-like cells sparsely distributed along the AP axis of the neural tube (**Fig. S12f**). These results suggest that human RA-gastruloids may recapitulate aspects of early neurogenesis.

We also sought to perform a more detailed analysis of somites, by reanalyzing paraxial mesoderm derivatives (*i.e.* somites, differentiation front, and myocytes) including the differential expression of spatial markers of somites’ rostrocaudal (*UNCX*; *TBX3*) and dorsoventral (*PAX3*; *PAX1*) organization (**Fig. S13**; **Table S5**). Whereas *PAX3* (dorsal) was strongly expressed in somites, *PAX1* (ventral) was not (**Fig. S13d-e**), suggesting that similar to the neural tube, somites in these human RA-gastruloids are dorsally biased, again presumably due to the lack of a Sonic hedgehog (SHH)-secreting notochord. Additionally, as seen in both vertebrate embryos^45^, as well as in the mouse gastruloids and mouse TLS models^2,15^, both *UNCX* (caudal) and *TBX18* (rostral) are expressed in a mutually exclusive manner (**Fig. S13c**), suggesting that we are reconstituting the rostral-caudal axis of somites. Consistent with this, we identify subsets of somitic cells that may correspond to migratory muscle precursors (*PAX7+*; *MET*+), myotome (*MYF5*+; *MET*+), syndetome (*SCX*+), sclerotome (*PAX9*+, *NKX3-2*+), endotome (*KDR*+) and an unknown cell type (*DNAJC5B*+) (**Fig. S13d-e**; **Table S5**).

Overall, these analyses and immunostaining experiments confirm that we are inducing neural tube and segmented somite-like structures in human gastruloids through the introduction of a discontinuous regime of retinoic acid. Although the dorsal-ventral axis is not fully established due to the lack of a notochord-equivalent SHH-secreting structure, human RA-gastruloids contain subpopulations of cells that resemble the more advanced cell types than previously achieved in either human or mouse gastruloid models, suggestive of operational signaling gradients along both the dorsoventral (neural crest) and mediolateral (intermediate mesoderm) axes. Particularly in the 120 hr gastruloids, we also observe the progression towards even more differentiated cell types (*e.g.* neural progenitors, cardiomyocytes, myocytes, renal epithelium), as well as the appearance of endothelium.

### Computational staging of human and mouse gastruloid models

The original report of human gastruloids assessed, based on morphological features and Tomo-seq^17^, that they model late CS8 to early CS9 in human development^1^, which corresponds roughly to embryonic day (E) 7.5 to E8.0 of mouse development. Because we observed more advanced cell types than have previously been reported in human or mouse gastruloids (*e.g.* neural crest, intermediate mesoderm, skeletal muscle, etc.), we hypothesized that the RA-gastruloids might model more advanced stages of human development. Further suggestive of this possibility, CS11 embryos have 13-20 pairs of somites, together with sclerotomes forming in the ventromedial part of the somites (**Table S6**), while in 120 hr human RA-gastruloids, we observe at least 12 pairs of somites (**Fig. 4a**). Since human gastruloids represent only the post-occipital part of human embryos, and also considering that human embryos have 4 pairs of occipital somites^46^, somite staging would place the 120 hr human RA-gastruloids as having reached roughly the equivalent of CS11. Moreover, we detect *PAX9*+, *NKX3-2*+ cells in sc-RNA-seq data, implying the onset of sclerotome formation (**Fig. S13d-e**). Taken together, these analyses suggest 120 hr human RA-gastruloids may model aspects of CS11 human embryos, which corresponds roughly to E8.75 to E9.5 in mouse development (**Table S6**).

**Figure 4.**
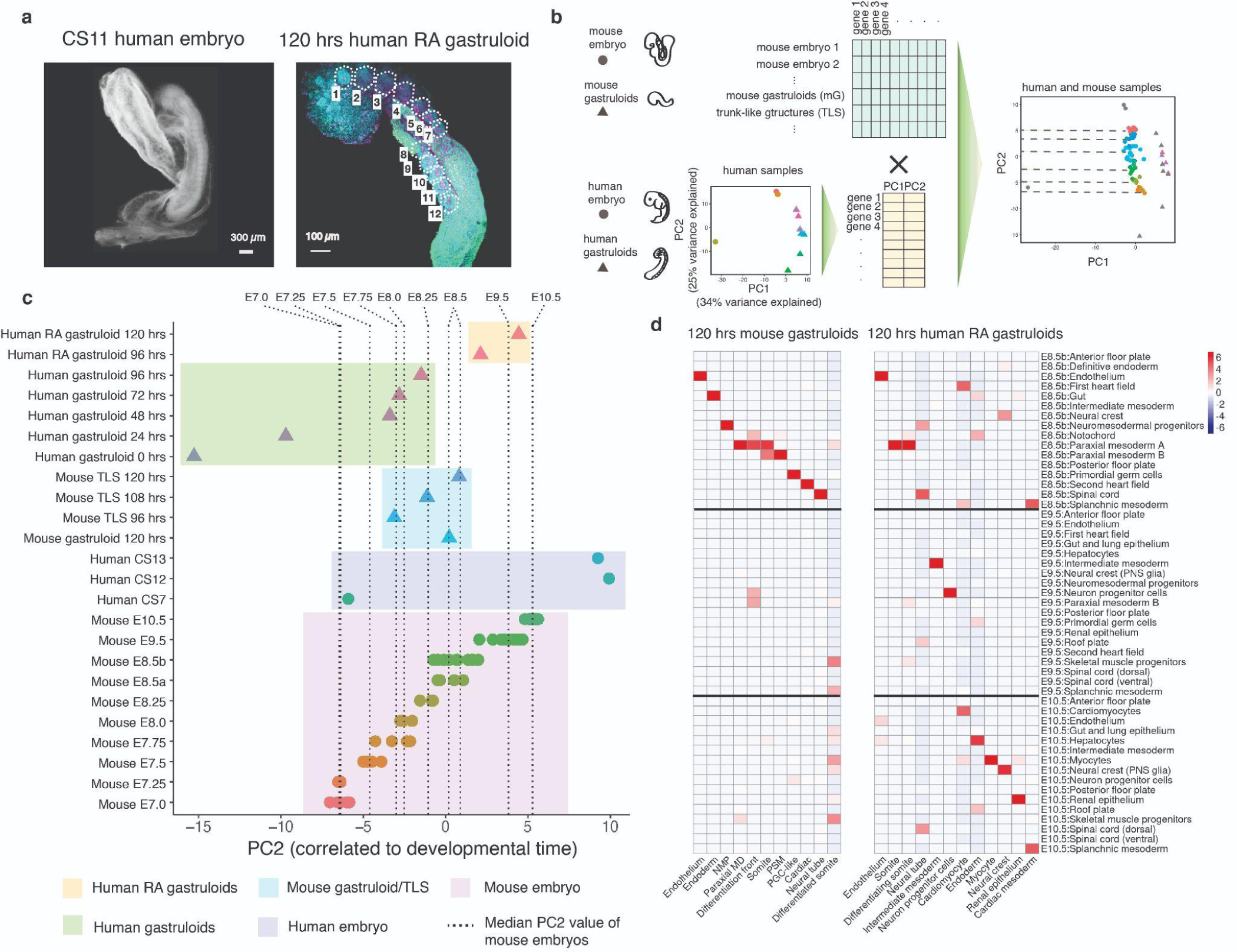
Computational staging of human RA-gastruloids and other mammalian synthetic embryo models. **a**, Morphology of human embryos at CS11 (left) and human RA-gastruloids at 120 hrs (right). **b**, Schematic of strategy for computational staging. In brief, PCA on human samples defines a principal component (PC) correlated with developmental progression (PC2). Projection of data from mouse samples onto this PC defined from human samples enables staging of the relative progression of synthetic embryo models across species and systems. **c**, Pseudo-bulk transcriptomes of pooled human RA-gastruloids (96 and 120 hrs), pooled original human gastruloids (0, 24, 48, 72 and 96 hrs), pooled mouse gastruloids (120 hrs)^2^, and pooled TLS (96, 108 and 120 hrs)^15^, individual human embryos at CS7, CS12, and CS13^18,47^, and individual^49,50^ or pooled^51^ mouse embryos projected onto the developmentally informative PC2 obtained from analysis of human samples. For mouse embryos, E8.5 and earlier samples are pooled embryos processed by 10X Genomics sc-RNA-seq^51^, while E8.5b and later samples are individual mouse embryos processed by sci-RNA-seq3^50,52^. **d**, NNLS-based cross-matching of cell types observed in 120 hr mouse gastruloids (left) or human RA-gastruloids (right) against cell types observed in E8.5-E10.5 mouse embryos^49,50^. Cell types observed in E8.5-E10.5 mouse embryos were selected based on the co-embedding shown in **Fig. S16**. Heatmap shows the scaled correlation coefficient (z-score) across all cell states in E8.5-10.5 mouse embryos.

However, these and other approaches to staging are somewhat *ad hoc*. As *in vitro* models continue to proliferate, there is a need for more systematic approaches to calibrate their relative progression, in relation to *in vivo* development. To this end, we sought to develop a framework for ‘computational staging’ of the developmental progression of human RA-gastruloids relative to somite-resolved mouse embryos, early human embryos, and other gastruloid models.

As a first step, we performed principal components analysis (PCA) of pseudobulk RNA-seq profiles of pools of human gastruloids and RA-gastruloids (96-120 hrs) along with two available datasets of human embryos at CS7^18^, CS12 and CS13^47^. As human gastruloids are derived from embryonic stem cells which will only give rise to embryonic lineages, we excluded cells derived from extra-embryonic tissues from these analyses. After dimensionality reduction, the second principal component (PC2) appeared strongly correlated with developmental time (**Fig. S14a**). The genes most correlated with PC2 included pluripotency related genes (*e.g. POU5F1*), master regulators of development (*e.g. LIN28A*), molecular markers of heart and neural lineages (*e.g. HES1, ZIC1, TNNT2*), and Hox genes (*e.g. HOXC9, HOXC6, HOXB5*) (**Fig. S15a-b**). Gene Ontology (GO) enrichment found positively correlated genes enriched for biological processes such as anterior-posterior pattern specification, epithelial and mesenchyme development, and embryonic organ development. In contrast, genes negatively correlated with PC2 are enriched for various metabolic processes, cell movement, and toxicity response, potentially corresponding to early events in response to induction with CHIR (**Fig. S15c**).

Along PC2, human gastruloids were correctly ordered with respect to time (**Fig. S14a**), while CS7 human embryos were positioned between 24 and 48 hr human gastruloids. CS7 is characterized by a notochordal process and gastrulation node^48^. Consistent with this, human gastruloids at 24 hr exhibited the emergence of *NOTO*+ cells (**Fig. S2**). Human gastruloids at 48-96 hrs and human RA-gastruloids at 96-120 hrs were placed between CS7 and CS12-13 human embryos. The placement is consistent with the fact that at least RA-gastruloids exhibit somite formation, indicating they have passed the pre-somitic stage at CS8. On the other hand, we do not observe the atrioventricular canal, thoracolumbar region, or hepatic sinusoids, which are signature structures at CS12^48^.

We next sought to achieve finer staging, as well as to relate the maturation of human gastruloids to *in vivo* mouse embryonic development, where we have access to much denser temporal sampling. In particular, for mouse E7-E10.5, a developmental window roughly equivalent to human CS7-CS13 (**Table S6**), sc-RNA-seq data is available from several sources, staged in either 6 hr^19^, 24 hr^49^, or single somite^50^ increments. Once again excluding cells derived from extraembryonic lineages, we generated pseudobulk RNA-seq profiles for staged mouse embryos, and then projected these onto the PC space described above, generated from pseudo-bulked human embryos and human gastruloid data (**Fig. 4b**). Sc-RNA-seq data from mouse gastruloid and TLS models was similarly processed and projected.

Despite the species difference, the staged mouse embryos were very well-ordered along the human-defined PC2, consistent with this principal component capturing some general aspect of mammalian developmental progression (**Fig. 4c**; **Fig. S14b-c**). Although CS7 human embryos were aligned with E7.0 mouse embryos, consistent with previous stage assessments, CS12-CS13 embryos were notably placed beyond E10.5 mouse embryos (**Fig. 4c**; **Table S6**). In this same framework, human gastruloids^1^ at 72 hrs map to E7.75-E8 mouse embryos, corresponding to CS8-CS9, consistent with staging by original report^1^. Both mouse gastruloids^2^ and TLS^15^ models at 120 hrs map to E8.5 mouse embryos. Finally, human RA-gastruloids at 120 hrs map to E9.5-E10.5 mouse embryos, consistent with their having developed to a more advanced stage than previously reported human and mouse gastruloids.

Given that gastruloids model only some aspects of *in vivo* development, and additionally that cell types observed within one model might be asynchronous, we sought to extend this computational staging approach to the level of individual cell types. After restricting to mouse embryo cell types present at E8.5-E10.5 that appear to also be observed in either 120 hr mouse gastruloids or 120 hr human RA-gastruloids (**Fig. S16**), we applied a non-negative least squares (NNLS)-based strategy^49^ to cross-match *in vitro* and *in vivo* cell types to one another, while leaving *in vivo* cell types observed at sequential timepoints in mouse development separate from one another (**Fig. 4d**). The cell types observed in 120 hr mouse gastruloids were individually very well aligned to cell types observed in E8.5 mouse embryos, consistent with the global analysis (**Fig. 4c**). However, the cell types observed in 120 hr human RA-gastruloids were more heterogeneously aligned to *in vivo* mouse cell types, *e.g.* with somites mapping to E8.5, intermediate mesoderm to E9.5, and neural crest to E10.5 (**Fig. 4d**). Although this could be due to differences between human and mouse development, we note that a similar analysis of 96 hr RA vs. non-RA human gastruloids^1^ finds that this heterogeneous alignment is specific to RA-gastruloids, as individual cell types within conventional (non-RA) human gastruloids are much more consistently aligned to mouse E8.5 cell types (**Fig. S17**).

Overall, these results show that human RA-gastruloids mature to a more advanced stage than previously described mammalian gastruloid models, roughly approaching the equivalent of mouse E9.5-E10.5. However, this greater progression may be accompanied by desynchronization with *in vivo* development, presumably due to the coarser signaling environment, lack of extra-embryonic cues, and other factors.

### Perturbation of WNT and BMP signaling in human RA-gastruloids

We next sought to evaluate the potential of human RA-gastruloids to serve as a screenable system by chemically perturbing canonical developmental signaling pathways during their differentiation. WNT signaling is known to crucially contribute to somite formation^34^. Correspondingly, exogenous CHIR, an agonist of WNT signaling, induces an excess of somites in the mouse gastruloid/TLS model^15^. To explore the role of WNT signaling in human RA-gastruloids, we modified the protocol to add CHIR back into the system at 48 hrs (**Fig. S18a**). Consistent with observations in mouse models^15^, re-addition of CHIR resulted in an excess of somite-like structures along the entire anterior-posterior axis of the gastruloid (**Fig. S18b**). The result reinforces the paradigm that the role of WNT signaling of somite development and segmentation is broadly conserved in vertebrate embryogenesis, through to humans.

BMP signaling is known to play various important roles in lineage segregation and maintenance during early development. To perturb BMP signaling in human RA-gastruloids, we modified the protocol to add either LDN193189 (hereafter LDN; a BMP inhibitor) or BMP4 into the system at 48 hrs onwards (**Fig. 5a**). With LDN, the SOX2-mCit-positive neural tube structure was consistently extended further to the anterior side of the gastruloid. In contrast, the length of the neural tube-like structure was shorter in BMP4-treated gastruloids and confined to the posterior side (**Fig. 5b**). This result indicates that BMP signaling has negative effects on neural tube elongation, as reported in various vertebrate animal models^53^. To explore the effects of BMP inhibition on other cell types, we performed sc-RNA-seq on LDN-treated RA-gastruloids (**Fig. 5c-e**). Consistent with the excessive elongation of the neural tube, we detected vastly more SOX2+/TBXT+/NKX1-2+ NMPs and SOX2+/PAX6+/SOX1+ neural tube cells than in untreated RA-gastruloids (**Fig. 5f-h**; **Fig. S19a-b**). On the other hand, LDN treatment also resulted in a paucity of intermediate mesoderm, renal epithelium, cardiac cells, neural crest cells, myotome, myocyte, and endotome (**Fig. 5f-h; Fig. S19d-e**). Of note, these cell types exhibited expression of ID genes (direct targets of BMP-SMAD signaling) in untreated RA-gastruloids (**Fig. S19c**).

**Figure 5.**
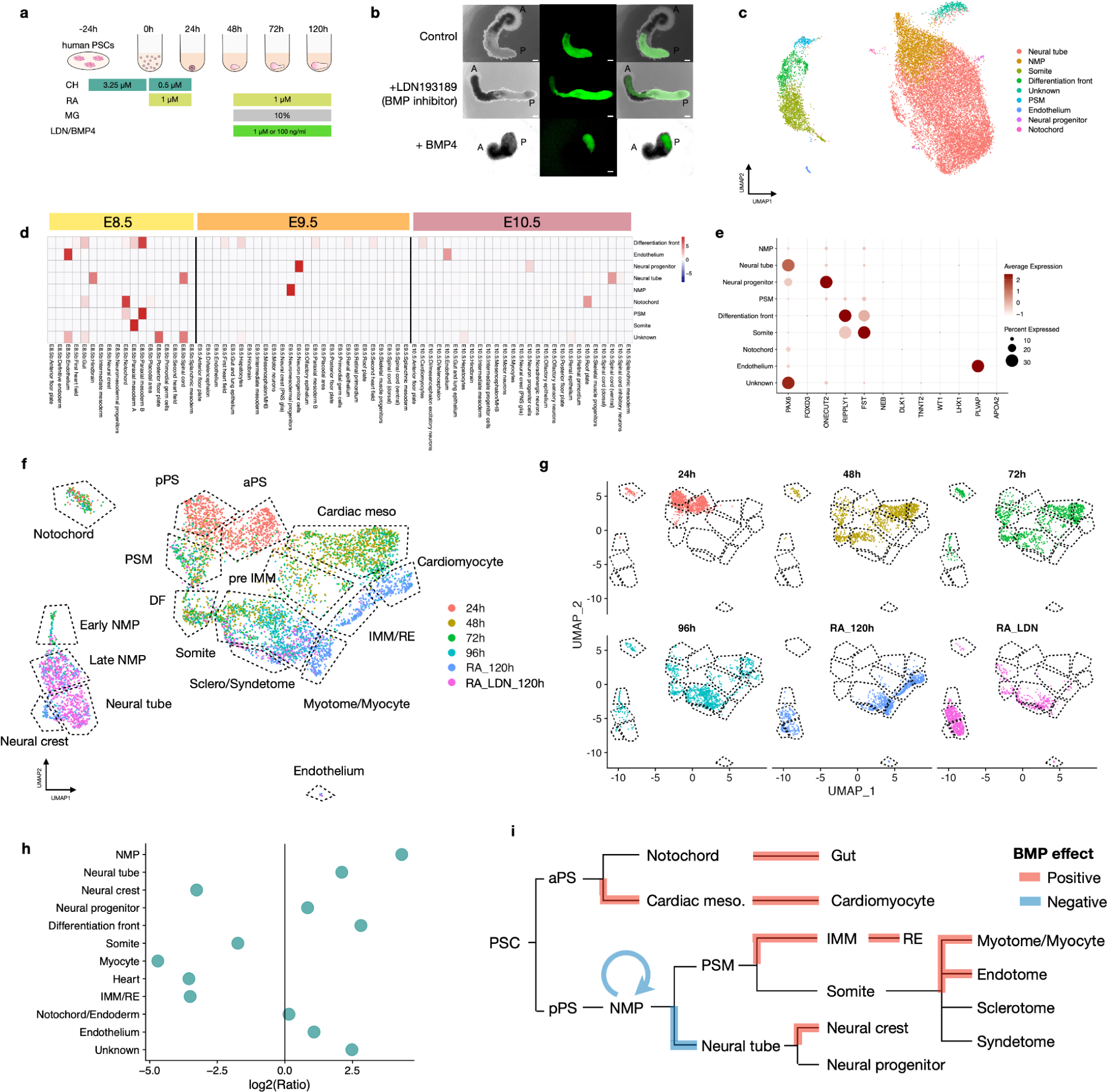
Effects of perturbing BMP signaling on the development of human RA-gastruloids. **a**, Schematic of perturbation of BMP signaling in human RA-gastruloids. CHIR, CHIR99021; LDN/BMP4, LDN193189 or BMP4. **b**, Representative images of wildtype (WT), LDN-treated, and BMP4-treated RA-gastruloids. A and P indicate the putative anterior and posterior of gastruloids, respectively. Scale bar, 100 µm. **c**, UMAP of sc-RNA-seq data derived from LDN-treated gastruloid. PSM, pre-somitic mesoderm; DF, differentiation front; NMP, neuromesodermal progenitor. **d**. NNLS-based correlation of cell types in LDN-treated RA human gastruloids (120 hrs) vs. cell types observed in E8.5-E10.5 mouse embryos. **e**, Bubble plot of marker gene expression in each cell type annotated in LDN-treated RA-gastruloid. **f**-**g**. Co-embedded UMAP of sc-RNA-seq data from non-RA-gastruloids (24-96 hrs), RA-gastruloids (120 hrs), and LDN-treated RA-gastruloids (120 hrs). 1,000 cells were used for each sample. aPS, anterior primitive streak; pPS, posterior primitive streak; PSM, pre-somitic mesoderm; DF, differentiation front; NMP, neuromesodermal progenitors; IMM, intermediate mesoderm; RE, renal epithelium. **h**, Cell type composition changes in LDN-treated RA-gastruloid, relative to untreated RA-gastruloids. **i**, Schematic view of the inferred effects of BMP signaling on the development of various cell types observed in human RA-gastruloids. PSC, pluripotent stem cell; aPS, anterior primitive streak; pPS, posterior primitive streak; NMP, neuromesodermal progenitor; PSM, pre-somitic mesoderm; IMM, intermediate mesoderm; RE, renal epithelium.

To examine the developmental progression of LDN-treated RA-gastruloids, we mapped them to *in vivo* mouse cell types^49^ from E8.5-E10.5 via NNLS cross-matching as described above. Interestingly, whereas NMPs from both the original gastruloids as well as RA-gastruloids are assigned to E8.5 (**Fig. 4d**, **Fig. S17**), NMPs in LDN-treated RA-gastruloids were assigned to E9.5 (**Fig. 5d**). A possible explanation is that in the untreated gastruloids, both the proliferation and maturation of NMPs is suppressed by BMPs, autonomously derived from intermediate mesoderm and cardiac cells (**Fig. S19b-c**). This is consistent with previous reports that BMPs may restrict the area within which NMPs can arise to the tailbud region^21,22,54^. However, in LDN-treated RA-gastruloids, BMP signaling is suppressed and this restriction removed, resulting in both proliferation and further maturation of the NMP pool in the posterior region, which contributes to a highly elongated neural tube.

However, although neural cells were greatly expanded, LDN-treated gastruloids were also markedly depleted for neural crest (**Fig. 5f-h**). This result can be explained through SOX2, a master regulator of neural stem cells^54^, constitutive expression of which inhibits neural differentiation^55^. BMP-mediated downregulation of *SOX2* is critical for both entering and exiting the neural stem cell state^53^, and indeed we observe *SOX2* downregulation (and upregulation of ID genes) in neural crest cells of untreated RA-gastruloids, whereas *SOX2* expression remains high (and neural crest cells essentially absent) in the neural tube cells of LDN-treated RA-gastruloids (**Fig. S19a-c**). A summary of inferred effects of BMP signaling on cell types observed in human RA-gastruloids is shown in **Fig. 5i**.

### Perturbation of key transcription factors in human RA-gastruloids

We next sought to investigate the potential for human RA-gastruloids as a screenable model for genetic perturbations. For this, we chose the transcription factors PAX3 and TBX6, since the corresponding mutants exhibit early lethality in mouse models^56,57^ and are therefore challenging to study *in vivo.* To knockout these genes, we introduced CRISPR-Cas9 RNA-protein complexes (RNPs) to human PSCs (**Fig. 6a**) and induced RA-gastruloids from these cells. While we didn’t detect morphological differences between PAX3-KO and non-targeting-control (NTC)-RNP RA-gastruloids, TBX6-KO RA-gastruloids showed a kinked posterior end with SOX2-mCit+ neural cells (**Fig. 6b**). On the other hand, marked differences were observed upon sc-RNA-seq profiling. PAX3-KO RA-gastruloids exhibited an increase in neural cells but a decrease in neural crest (**Fig. 6c-g**; **Fig. S20**), suggesting neural crest differentiation was inhibited by knockout of PAX3. This is consistent with a previous finding reporting dysregulation of neural crest cell development in *Pax3*-deficient or Splotch mutant mice^58,59^.

**Figure 6.**
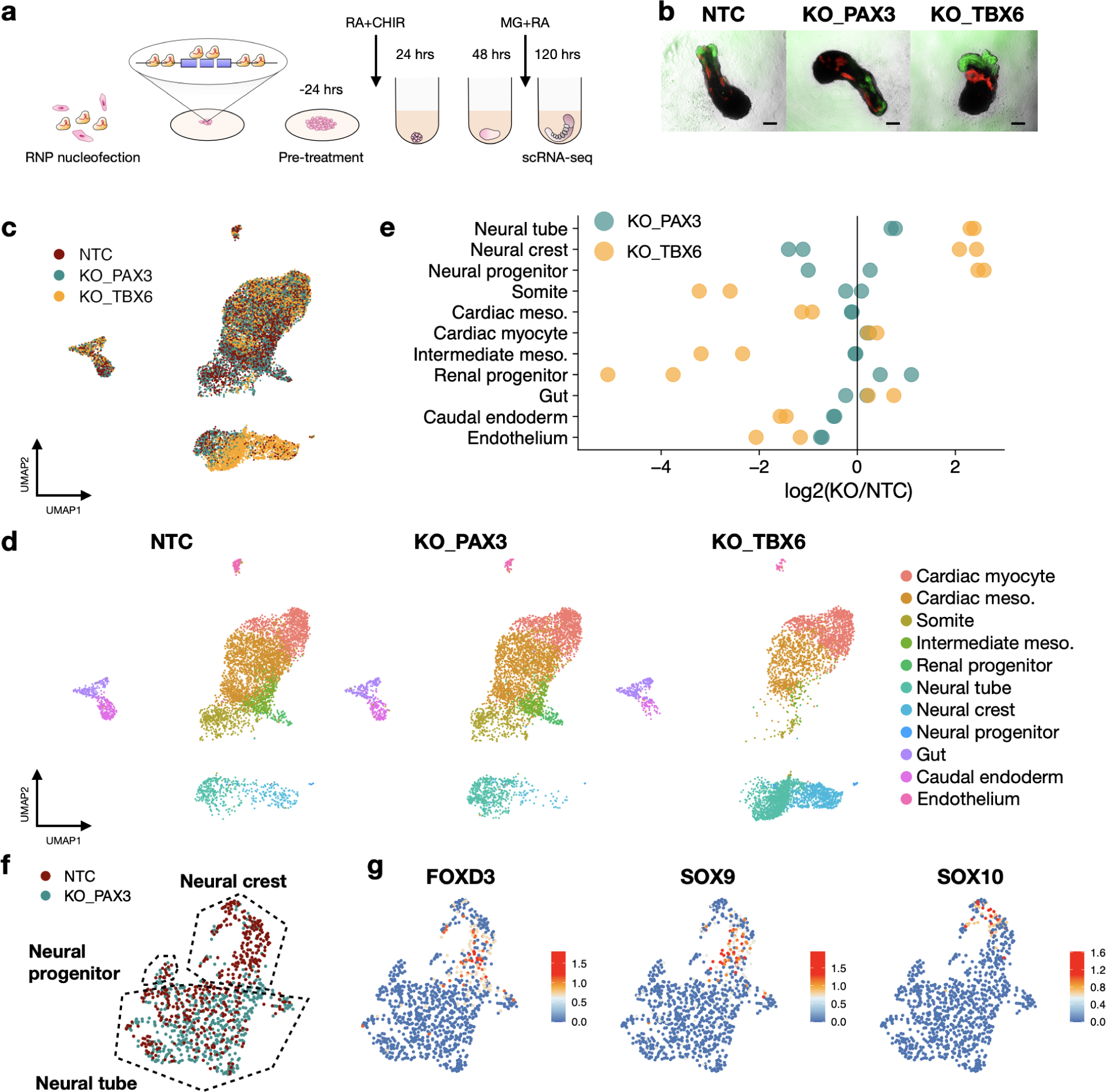
Genetic perturbation of transcription factors in human RA-gastruloids. **a**, Schematic of transcription factor knockouts in human RA-gastruloids using CRISPR/Cas9 RNPs. Six Cas9-gRNA RNPs were nucleofected into PSCs, inducing indels or full deletion of exons of *PAX3* or *TBX6*. Nucleofected PSCs were subjected to RA-gastruloid induction. RA-gastuloids were collected at 120 hrs after cell aggregation. **b**, Representative images of non-targeting-control (NTC), PAX3-KO, TBX6-KO gastruloids. N = 24 gastruloids for each condition. **c-d**, UMAP of sc-RNA-seq data from NTC, PAX3-KO, and TBX6-KO gastruloids, labeled by genotype (**c**) or cell types (**d**). **e**, Cell type composition changes upon knockout of each transcription factor. Log2 (KO/NTC) fold changes for each cell type are shown as dots. Vertical black line corresponds to no change in proportion of the cell type between KO and NTCs. **f**, Sub-clustering analysis on neural cells (neural tube, neural crest, and neural progenitor cells). **g**, Expression of marker genes of neural crest cells.

In mice, Tbx6 is an established master regulator of somitic lineages differentiation and suppressor of neuronal lineages through the repression of a *Sox2* enhancer^57,60^. Consistent with observations in mice, we observed a markedly higher number of neural cells (including neural crest), in TBX6-KO human RA-gastruloids (**Fig. 6c-e**). In contrast, somite cells were almost absent. We also observed the loss of intermediate mesoderm and renal epithelium in TBX6-KO RA-gastruloids (**Fig. 6d**), suggesting that intermediate mesoderm and renal cells in human RA-gastruloids derive from TBX6-positive PSMs, as has been recently observed in mouse embryos^61,62^. Unexpectedly, we also found loss of a specific population of definitive endoderm cells (*CDX2*/*FGF17*+) in TBX6-KO RA-gastruloids (**Fig. 6d-e**; **Fig. S20**). This may be a secondary effect of the loss of PSM, which may contribute to the appropriate posterior environment for definitive endoderm in the tailbud region, but warrants further investigation.

Overall, we conclude that RA-gastruloids can serve as a versatile and robust model for investigating the cellular and molecular mechanisms by which cell types emerge in human post-implantation development, via both chemical and genetic perturbation.

## DISCUSSION

The field of synthetic mammalian embryology is on a remarkable trajectory. For both human and mouse, “stembryo” models open up early development to detailed observational and perturbational investigations that are simply not possible *in vivo*, for both practical and ethical reasons. In this study, we show that a specific, discontinuous regimen of retinoic acid (RA) robustly induces human gastruloids with embryo-like morphological structures, including a neural tube and segmented somites. Although simpler conditions have yielded concurrent somite and neural tube formation in mouse gastruloids^2,15^, to our knowledge, a human neural tube has not previously been induced *in vitro* from scaffold-free stem cell aggregates, let alone concurrent somitogenesis and neural tube formation.

It remains unclear why a discontinuous regimen of RA leads to formation of a neural tube and segmented somites in human gastruloids, while Matrigel is sufficient to achieve this in mouse gastruloids^2,15^. One possibility is that the balance between WNT and RA signaling that underlies the bi-potency of NMPs is considerably different in human gastruloids, but is restored by supplemental RA at specific intervals. In terms of explaining the more advanced cell types observed (neural crest cells, renal progenitor cells, skeletal muscle cells, and, rarely, neural progenitor cells), it is possible that RA directly contributes to induction of some of these (*e.g.* RA has been used for the induction of intermediate mesoderm from PSCs^63–66^). However, RA is not thought to contribute to the induction of neural crest^67–70^, so it is possible that their appearance is a secondary consequence of RA, *e.g.* following from the emergence of a dorsal neural tube.

The proliferation of stembryo models is uncovering some new challenges. For example, it is difficult to temporally calibrate these models against *in vivo* development, as well as against one another, based on purely morphological criteria. In particular, morphological features are not evident in some models^1^, and *in vitro* models may have different morphology than *in vivo* development for secondary reasons, *e.g.* different spatial constraints on their development. To address this, we sought to develop and apply a systematic approach to computationally stage *in vitro* gastruloid models relative to *in vivo* timepoints based on sc-RNA-seq data. The approach facilitates temporal alignment of data from human and mouse, while also highlighting asynchronous developmental progression within models, as has been observed in both normal and disease-modeling organoids^71–73^. Looking forward, we anticipate that the further application of this approach will facilitate the systematic delineation of which specific aspects of development (with respect to both cell type(s) and time-point) are modeled by each of the expanding number of stembryo and organoid systems. For example, while this manuscript was in preparation, three papers or preprints appeared describing hPSC-derived “somitoid/axioloid” models^16,74,75^. We evaluated sc-RNA-seq data from one of these studies^16^ with our computational staging method (**Fig. S21a-c**). The 168 hrs human somitoid mapped to the E8.5 stage of mouse development (**Fig. S21d**), with, as reported, cell types limited to mainly somitic lineages (**Fig. S21b**).

Increasingly sophisticated stembryo models may also raise ethical concerns that might be mitigated through computational staging. For example, concerns about neural activity have been raised in the context of brain organoids^76,77^, while blastoids^8,78,79^, which contain all extraembryonic tissues, could at least theoretically develop into advanced embryos. Although the appearance of the primitive streak was used to establish the “14 day rule” for *in vitro* culture of fertilized human eggs^80^, this rule makes little sense for stembryo and organoid models, particularly as they might mature slower or faster than *in vivo* development. We propose that computational staging represents a more systematic and universal approach for assessing the developmental progression and completeness (or lack thereof) of complex *in vitro* models of development. For example, here we estimate ∼5 day old human RA-gastruloids to have matured to the equivalent of ∼29 day old human embryos (CS11), but also unequivocally confirm that they lack anterior neural structures as well as extraembryonic tissues.

The efficiency with which embryo-like morphological structures form in human RA-gastruloids is much greater than reported in related mouse systems (89% in human RA-gastruloids [both neural tube and segmented somites] vs. 50% in mouse TLS^15^ [segmented somites] vs. 4% in Matrigel-embedded mouse gastruloids^2^ [striped Uncx4.1 expression]). This robustness makes RA-gastruloids an attractive model for detailed investigations of early human development. It is particularly important to arrayed screens, in which substantial technical variation between wells would markedly compromise statistical power. Here we illustrate this potential via a limited number of chemical and genetic perturbations, but we envision tremendous possibilities for further scaling of human RA-gastruloids as a simple, robust model for decoding many aspects of early human embryogenesis, as well as for certain classes of human developmental disorders (*e.g.* neural tube defects).

## MATERIALS & METHODS

### Human ES cell culture

The pluripotent stem cell lines, hESCs (RUES2-GLR), were gifted by Dr. Ali Brivanlou (Rockefeller University). Human ES cells were maintained in StemFlex medium (Thermo, A3349401) on Geltrex (Thermo, A1413201) and were routinely passaged using StemPro Accutase (Thermo, A1110501) to new Geltrex-coated wells as recommended by the manufacturer. For the first 24 hrs after passaging, hESCs were cultured in StemFlex with 10 μΜ Y-27632 (Sellek, S1049) to prevent apoptosis^81^.

### Human RA-gastruloid induction

4×10^4 hESCs were seeded on 0.5 μg/cm^2^ Vitronectin-coated 12-well plate (Gibco, A14700) in Nutristem hPSC XF medium (Biological Industries, 05-100-1A) with 10 µM Y-27632. On day 1, the medium was replaced with Nutristem containing 5 µM Y-27632. On day 2, the medium was replaced with Nutristem. On day 3, the medium was replaced with Nutristem containing 3.25 µM CHIR (Millipore, SML1046). 24 hrs later, cells were detached by StemPro Accutase and dissociated into single cells. 4,000-5,000 cells were seeded to 96 wells with 50 µl Essential 6 medium (Thermo, A1516401) containing 0.5 µM CHIR, 0.5 or 1 µM RA (Millipore Sigma, R2625), and 5 µM Y-27632. At 24 hrs, 150 µl of Essential 6 medium was added to each well. At 48 hrs, 150 µl of the medium was removed with a multi-channel pipette, and 150 µl of Essential 6 medium containing 5% Matrigel and 1 µM RA was added and maintained in 37C 5% CO2 incubator until 120 hrs. To further test the effect of the signal perturbation on gastruloid formation, we added 1 uM LDN-193189 (STEMCELL Technologies, 72147), 100 ng/ml BMP4 (R&D, 314-BP-010), or 3 uM CHIR to the medium at 48 hrs along with 5% Matrigel and 0.5 µM RA.

### Genetic perturbation with CRISPR/Cas9 RNPs

RNP complexes were prepared according to the manufacturer’s procedures. Briefly, equal molars of crRNA and tracrRNA (IDT, 1072532, listed in **Table 7**) were mixed and heated at 95C for 5 minutes in a thermal cycler and kept at room temperature for 10-20 min for hybridization. AltR-Cas9 protein (IDT, 1081058) was added to the crRNA and tracrRNA mixture to assemble Cas9-RNPs. RUES2-GLR hESCs were trypsinized with StemPro Accutase and the reaction was quenched with StemFlex media supplemented with 10 mM Y-276322. 2×10^5 cells were transferred to a new tube and centrifuged at 250g for 5 min. Cells were resuspended in 20 µl nucleofection buffer (16.4 µl Nucleofector® Solution + 3.6 µl Supplement) provided in P3 Primary Cell 4D-Nucleofector™ X Kit S (Lonza, V4XP-3032). After the addition of 3 µl RNP and 0.5 µl of Alt-R Cas9 Electroporation Enhancer (IDT, 1075915), cells were transferred to 16-well Nucleocuvette™ Strips and nucleofected with the CA-137 program. The nucleofected cells were transferred to the 12-well plate which contains Nutristem with 10 mM Y-27632 and after 24 hrs, the medium was replaced with Nutristem without Y-27632. Cells were maintained until they reach 50-70% confluent. Then, RNP-introduced cells were transferred onto 0.5 μg/cm^2^ Vitronectin-coated 12-well plate and proceeded to RA-gastruloid induction steps described above and collected at 120 hrs of induction for sc-RNA-seq analysis.

### Immunostaining of gastruloids

For whole-mount immunostaining, gastruloids were fixed in 4% paraformaldehyde overnight at 4 °C, washed with PBST (0.2% Tween20), soaked in blocking buffer (PBS containing 0.1% BSA and 0.3% Triton X-100) overnight at 4 °C and then incubated with primary antibodies diluted with blocking buffer overnight at 4 °C. The samples were washed with washing buffer (PBS containing 0.3% Triton X-100), incubated with secondary antibodies and DAPI overnight at 4 °C, and washed and mounted in Fluoro-KEEPER antifade reagent (Nacalai USA). For phalloidin staining, Alexa Fluor™ 647 Phalloidin (A22287; Thermo) was added to the secondary antibody at a dilution of 1:400. All samples were analyzed with an LSM710 (Zeiss) or LEICA SP8X (Leica) confocal microscope with negative control samples where the primary antibodies were not added. The antibodies used in this study are listed in **Table S8**.

### Single cell-RNA-seq of gastruloids

Gastruloids (n = 48-96) were collected from 96 wells using a micropipette and transferred to PBS (-) to wash out the medium and incubated with 0.05% Trypsin-EDTA for 8 min at 37C. After quenching the reaction by the addition of 1/10 volume of FBS, gastruloids were dissociated into single cells by repeated pipetting. Dissociated single-cell suspensions were loaded in Chromium Next GEM Chip G Single Cell Kit (Chromium, PN-1000120) on a 10X Chromium controller according to the manufacturer’s instruction. Single-cell RNA-seq libraries were generated with Chromium Next GEM Single Cell 3’ GEM, Library & Gel Bead Kit v3.1 (Chromium, PN-1000121) and Dual Index Kit TT Set A, 96 rxns (Chromium, PN-1000215). Library concentrations were determined by Qubit (Invitrogen) and the libraries were visualized by Tapestation (Agilent). All libraries were sequenced on NextSeq 2000 (Illumina) (Read 1: 28 cycles, Read 2: 85 cycles, Index 1: 10 cycles, Index 2: 10 cycles).

### Processing of sequencing reads

Base calls were converted to fastq format using cellranger v6.0.0^82^ mkfastq function. The sequencing reads are then demultiplexed based on i5 and i7 barcodes, mapped to hg38 reference genome and assigned to GRCh38 (GENCODE v32/Ensembl 98) genes by cellranger v6.0.0 count function with default setting.

### sc-RNA-seq analysis

The filtered feature barcode matrices generated by cellranger were then loaded on R and converted to Seurat objects with Seurat v4.1.1^83^. The threshold for UMI counts and feature counts were determined by the valley in bimodal distribution of UMI counts and feature counts respectively. Cells showing upper outlier values in the violin plot of UMI counts and feature counts are also excluded to remove potential doublets. To further make sure all doublets are removed, Scrublet was used to calculate a doublet score for each cell. Cells with doublet scores higher than the simulated threshold value by Scrublet are potential doublets with two different cell types and they are excluded from the dataset. Since we have observed strong cell cycle effects in our dataset after initial dimensional reduction, we decided to remove cell-cycle related genes^84^ (specifically, genes with prefix HIST, MT, TOP, CDK, CCN, CDC, CCDC, MKI; genes containing MALAT, AUR and genes with suffix NUSAP, SMC, CENP, UBE, SGO, ASPM, PLK, KPN, RP, PTTG, SNHG, CK, BUB, KIF, KCNQ, SMO, HMG, S100, LINC, ATP, IGFBP, HSP, FOS, JUN) were removed from the datasets. After quality control, each dataset was log-normalized and regressed out heterogeneity associated with cell cycle stage and the mitochondrial contamination by SCTranform. The top 3000 variable genes identified by SCTranfom were then used for dimensionality reduction by PCA. Then the first 30 PCs were used for dimensionality reduction by UMAP. Cells were clustered by graph-based clustering algorithm in which first cells were first embedded in a K-nearest neighbor based graph structure and then Louvain algorithm was applied to detect cell clusters.

Time-course sc-RNA-seq datasets were integrated by Harmony implemented in Seurat. Briefly, datasets from each timepoint were merged together. The merged dataset was log-normalized and the top 2,000 variable genes were selected for downstream integration. Then, the merged dataset was scaled and heterogeneity associated with cell cycle stage and the mitochondrial contamination were regressed out. Dimensionality reduction by PCA was implemented to generate a low-dimensional embedding of cells in the merged dataset. The low-dimensional embedding was then input into Harmony to correct for batch effects among samples collected from different experiments.

To compare gastruloid cell types and embryo cell types in the Mouse Organogenesis Cell Atlas (MOCA) as well as data more recently generated from E8.5 via sci-RNA-seq3^49,50^, 50,000 mouse embryonic cells were randomly selected at each timepoint spanning E8.5-13.5 from MOCA (28 - 19694 cells representing each cell type) and all cells from human RA-gastruloids (120 hrs) and mouse gastruloids (120 hrs) for integration. A lift-over list of mouse and human genes was downloaded from BioMart-Ensembl (Ensembl Genes 102). The gene features in mouse and human datasets were converted to a list of human and mouse orthologous pairs. The relationship of multiple homologs to one gene in the other species were retained in this case. Human gastruloids and mouse gastruloids datasets were integrated with mouse embryo data with the Seurat V3 integration algorithm^85^.

### Global staging of *in vitro* models

10x Genomics sc-RNA-seq data from CS12-16 human embryos^47^, Smart-seq2 sc-RNA-seq data from a CS7 human embryo^18^, 10x Genomics sc-RNA-seq data from mouse embryos E6.5-8.5^51^, sci-RNA-seq data from mouse embryos E8.5-13.5^49,50^, 10x Genomics sc-RNA-seq data from mouse gastruloids^2^ and TLS^15^ were collected online. Time-course sc-RNA-seq datasets and 96 hr and 120 hr RA-gastruloid datasets after pre-processing were used for staging. Cells with transcriptomes mapping to extraembryonic tissues were excluded from all embryo datasets. For CS12-16 human embryos, hemoglobin, mitochondrial, sex-specific, cell cycle, and batch-effect genes were removed following methods from the original paper. For the CS7 embryo, cell cycle and mitochondrial genes were removed. Since CS7 was sequenced using Smart-seq2 which has more variations in library depth between single cells and uses read counts instead of UMI counts, additional normalization with Seurat V4.1^83^ NormalizeData function was performed on the CS7 dataset. After normalization, the read distribution for each cell approximated a Poisson distribution with a similar library size range as UMI counts from other datasets.

Where possible, all embryo datasets were pseudobulked by embryo. Specifically, all UMI counts connected to a particular embryo sample (or embryo sample, in the case of pools) were added together for each gene. Human and mouse gastruloids were pseudobulked by sample (as these derive from multiple embryos). Specifically, all UMI counts connected to a particular gastruloid sample were added together for each gene.

Since we are mainly focusing on human early embryonic development, highly variable genes from human embryo developmental datasets^18,47^ were used for down-stream analysis. For CS12-16 embryos, highly variable genes were calculated based on Poisson distribution^86^ and a total of 1,361 HVGs were obtained. We then performed intersection of embryo CS12-16 HVGs, features in CS7 embryos, human gastruloid samples and mouse gastruloid samples were taken for future co-embedding, after converting the gene features to mouse human orthologous pairs as described above. The intersection resulted in a total of 455 genes for PCA analysis.

The pseudobulk profiles of human embryos CS12-13, human gastruloids (0-96 hrs) and human RA-gastruloids (96-120 hrs) were used to construct a PCA space with the 455 genes. Specifically, the pseudobulk matrix was normalized and scaled by Seurat V4.1^83^. PCA analysis was performed on the scaled matrix. The scaled expression of genes in each sample were subjected to Pearson’s correlation with the PC2 embeddings of the sample in order to obtain top genes highly correlated/anti-correlated with PC2. Mouse embryos and mouse gastruloids were projected onto the human PCA space by multiplying the loadings of the human PCA space to the scaled matrix of human and mouse gastruloids, human and mouse embryos.

### Lineage specific staging of *in vitro* models

Expression values for each gene in each embryonic cell type were calculated by summing the log-transformed normalized UMI counts of all cells of that type across all embryos in each time point from E8.5 to E10.5. In order to minimize batch effects, only mouse embryos prepared by sci-RNA-seq were selected^49,50^. Only cell types showing mapping with cell types in human or mouse gastruloids in previous co-embedding were included for the lineage specific staging. Expression values for each gene in each annotated gastruloid cell type were calculated by summing the log-transformed normalized UMI counts of all cells of that type in each gastruloid sample.

To compare cell types in gastruloids toward selected cell types in mouse embryos (E8.5-E10.5), homologous genes were taken on both datasets and nonnegative least-squares regression were applied to predict gene expression in target cell type (Ta) in dataset A based on the gene expression of all cell types (Mb) in dataset B, Ta = β0a + β1aMb, based on the union of the 200 most highly expressed genes and 200 most highly specific genes in the target cell type. Datasets A and B were then switched for prediction; that is, predicting the gene expression of target cell type (Tb) in dataset B from the gene expression of all cell types (Ma) in dataset A, Tb = β0b + β1bMa. Finally, for each cell type a in dataset A and each cell type b in dataset B, two correlation coefficients, β = 2(βab + 0.001)(βba + 0.001), were combined to obtain a statistic for which high values reflect reciprocal, specific predictivity. The β score were then scaled across all selected cell types across E8.5-E10.5. The stage for each cell type in gastruloids was selected by identifying embryonic cell types with a scaled β score greater than 5 and a 1-to-1 relationship or averaging the developmental time between 2 cell types with the highest scaled scores.

## Ethics Statement

Our research was subject to review and approval from the Embryonic Stem Cell Research Oversight (ESCRO) of the University of Washington.

## Acknowledgments

We thank the members of the Shendure Lab, as well as members of the Allen Discovery Center for Cell Lineage Tracing, for helpful feedback and discussions; A. Boulgakov for helping with cell culture experiments; A.M. Arias, N. Moris, A. Alemany and SC van den Brink for helpful discussions; Dr. Ali H. Brivanlou for providing RUES2-GLR ESCs; the TOM & SUE ELLISON STEM CELL CORE of Institute for Stem Cell and Regenerative Medicine (ISCRM), and KECK microscopy center of University of Washington. This work was supported by a grant from the Paul G. Allen Frontiers Group (Allen Discovery Center for Cell Lineage Tracing to JS) and the National Human Genome Research Institute (UM1HG011586 to JS). JS is an Investigator of the Howard Hughes Medical Institute.

## Author Contributions

N.H. and J.S. designed the research. W.Y. C.K. and N.H performed experiments. W.Y. and N.H. performed computational analysis. C.K., C.Q., S.S. assisted with data analysis. B.K.M., R.K.G., S.G.R., C.L., R.M.D., and S.S. assisted with the interpretation of results. E.N. assisted with the imaging of gastruloids. J.S., W.Y. and N.H. wrote the paper, with input from all authors.

---

**Table S1.**
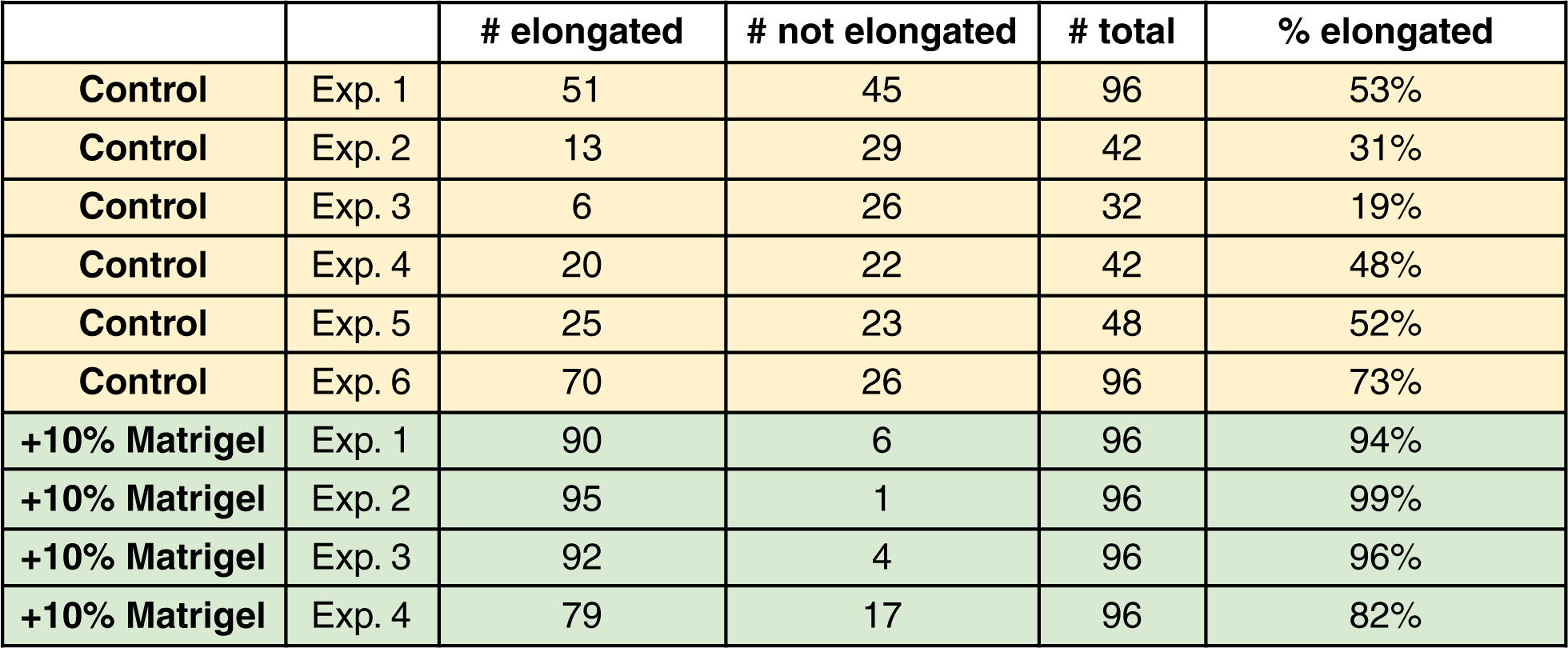
Ratio of elongated human gastruloids with the addition of Matrigel. 10% Matrigel was added at 48 hrs after induction.

|  |  | # elongated | # not elongated | # total | % elongated |
| --- | --- | --- | --- | --- | --- |
| **Control** | Exp. 1 | 51 | 45 | 96 | 53% |
| **Control** | Exp. 2 | 13 | 29 | 42 | 31% |
| **Control** | Exp. 3 | 6 | 26 | 32 | 19% |
| **Control** | Exp. 4 | 20 | 22 | 42 | 48% |
| **Control** | Exp. 5 | 25 | 23 | 48 | 52% |
| **Control** | Exp. 6 | 70 | 26 | 96 | 73% |
| **+10% Matrigel** | Exp. 1 | 90 | 6 | 96 | 94% |
| **+10% Matrigel** | Exp. 2 | 95 | 1 | 96 | 99% |
| **+10% Matrigel** | Exp. 3 | 92 | 4 | 96 | 96% |
| **+10% Matrigel** | Exp. 4 | 79 | 17 | 96 | 82% |

**Table S2.** Cell type proportions in human gastruloids (96 hrs), mouse gastruloids (120 hrs), and mouse TLS (120 hrs). N.D. = Not detected.

| (%) | Human gastruloids<br>@ 96 hrs | Mouse gastruloids<br>@ 120 hrs | Mouse TLS<br>@ 120 hrs |
| --- | --- | --- | --- |
| NMP | 4.5 | 13.3 | 10.6 |
| Neural Tube | *N.D.* | 13.5 | 28.1 |
| PSM | 9.2 | 13.4 | 1.9 |
| Differentiation front | 9.0 | 9.2 | 3.1 |
| Somite | 50.5 | 41.2 | 48.5 |
| Advanced Mesoderm | 6.9 | 1.4 | 2.5 |
| Cardiac | 13.8 | 4.0 | *N.D.* |
| Node | 3.9 | *N.D.* | *N.D.* |
| Gut | 2.3 | 0.4 | 1.5 |
| Endothelium | *N.D.* | 2.3 | 2.7 |
| PGCLC | *N.D.* | 1.2 | 1.1 |

**Table S3. Differentially expressed genes for annotated cell type clusters at 24, 48, 72, and 96 hrs after gastruloid induction**. Default thresholds from the FindAllMarkers function of the *Seurat* V4.0.1 package were used.

**Table S4. Marker genes for neural sub-clusters.** Default thresholds from the FindAllMarkers function of the *Seurat* V4.0.1 package were used to determine these marker genes.

**Table S5. Marker genes for somite sub-clusters.** Default thresholds from the FindAllMarkers function of the *Seurat* V4.0.1 package were used to determine these marker genes.

**Table S6. Comparison of human vs. mouse in vivo developmental stages.** Summary of characteristic features (based on citations) at an aligned set of human and mouse early developmental stages^49,55,56^.

**Table S7. List of crRNAs used in this study.**

**Table S8. List of antibodies used in this study.**

**Figure S1.**
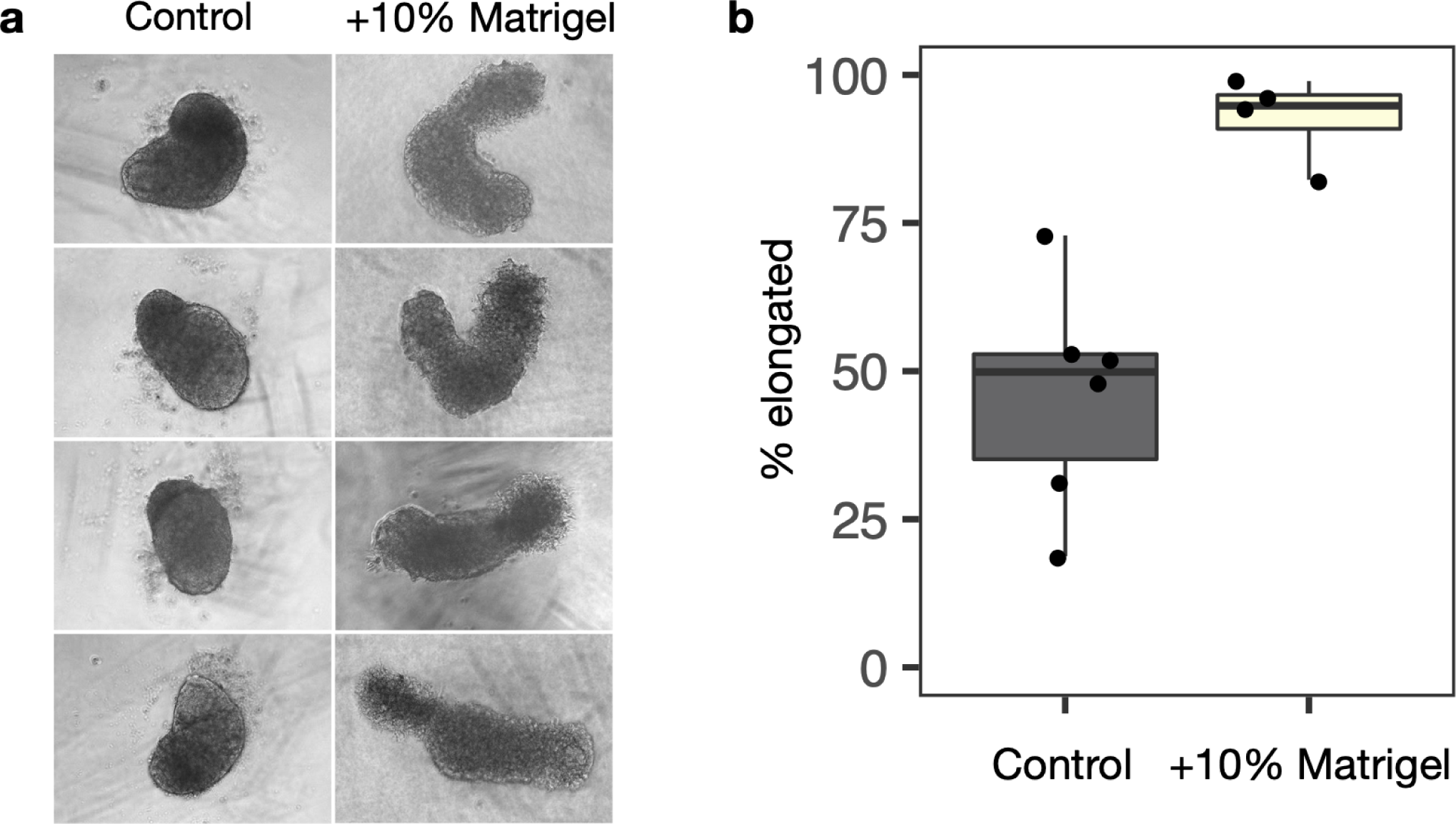
Effects of Matrigel on human gastruloid morphology. **a**, Representative images of human gastruloids without (left) vs. with (right) 10% Matrigel. Note that addition of 10% Matrigel enhances gastruloid elongation. **b**, Boxplot showing proportion of elongated gastruloids observed in the absence (left) vs. presence (right) of 10% Matrigel across a total of ten experiments. Raw counts are provided in **Table S1.**

**Figure S2.**
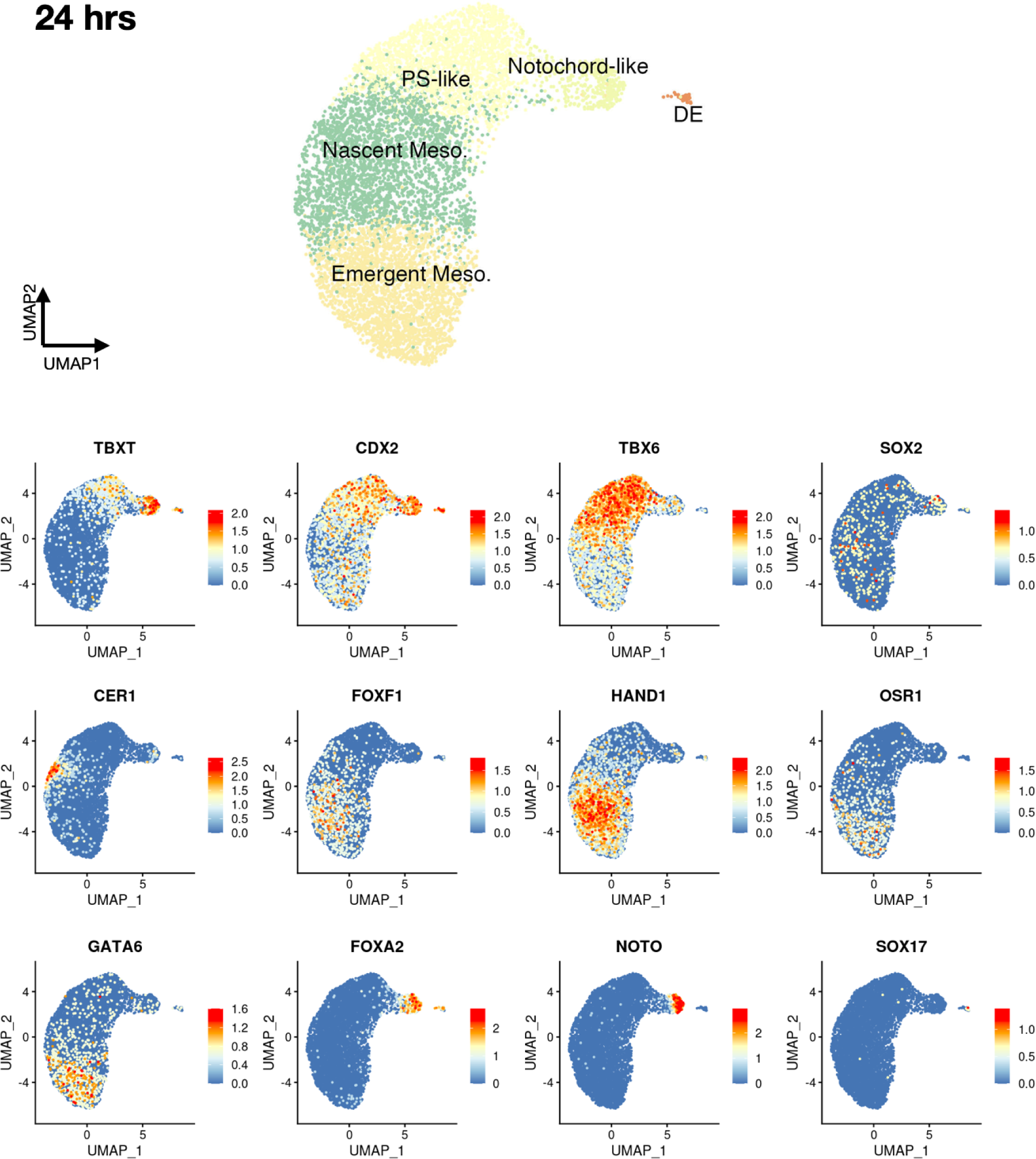
Marker gene expression and cell type annotation in human gastruloids at 24 hrs after induction. UMAP with unsupervised clustering of sc-RNA-seq profiles is shown, along with expression of marker genes used to annotate these clusters as a continuum of primitive streak-like, nascent mesoderm and emergent mesoderm, along with notochord-like cells and definitive endoderm. DE, definitive endoderm; PS, primitive streak.

**Figure S3.**
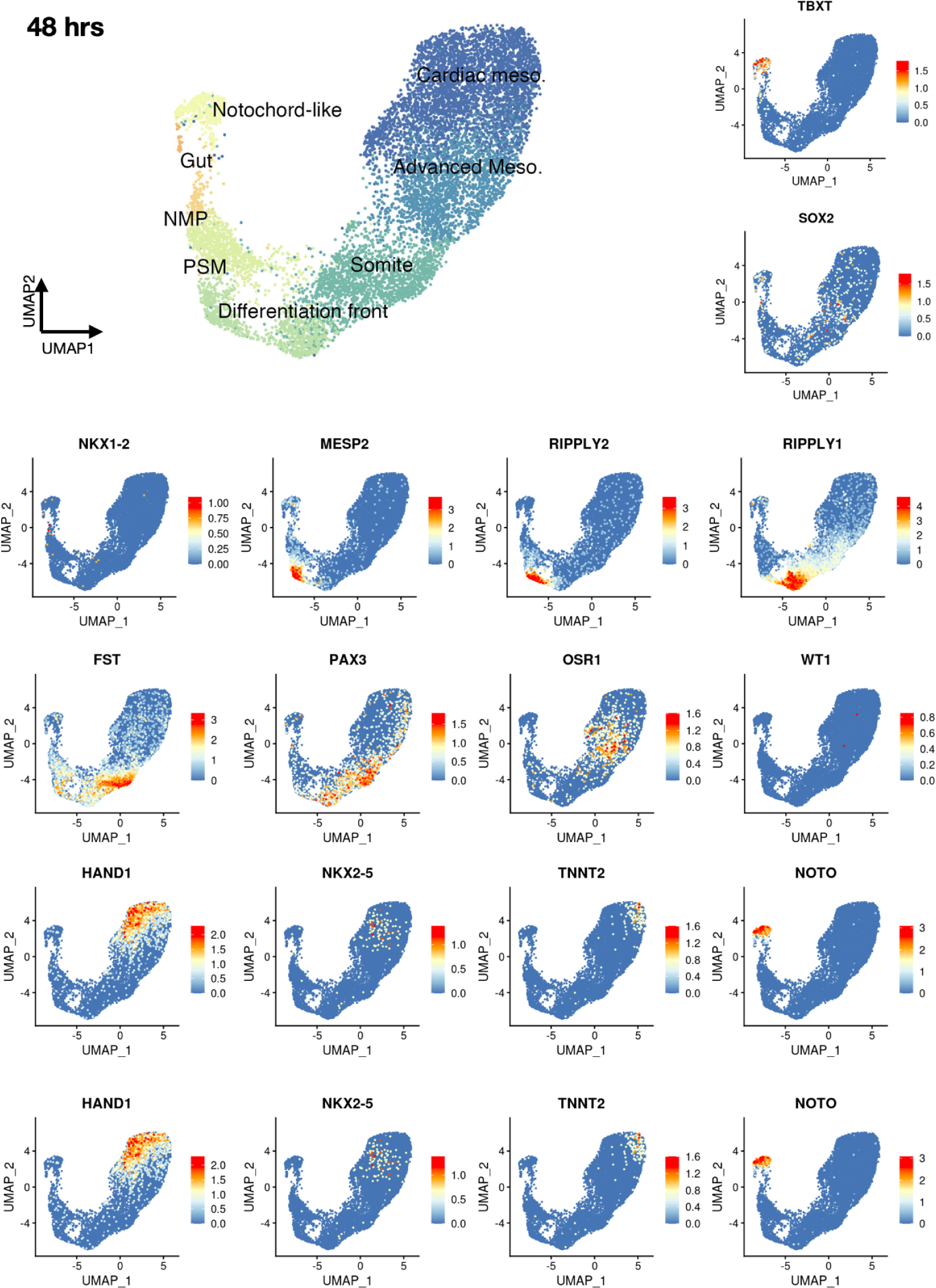
Marker gene expression and cell type annotation in human gastruloids at 48 hrs after induction. UMAP with unsupervised clustering of sc-RNA-seq profiles is shown, along with expression of marker genes used to annotate these clusters as a continuum of neuromesodermal progenitors, presomitic mesoderm, differentiation front, and somites; additionally, advanced mesoderm and cardiac mesoderm, as well as notochord-like and gut/endoderm-like cells. NMP, neural mesodermal progenitor; PSM, presomitic mesoderm.

**Figure S4.**
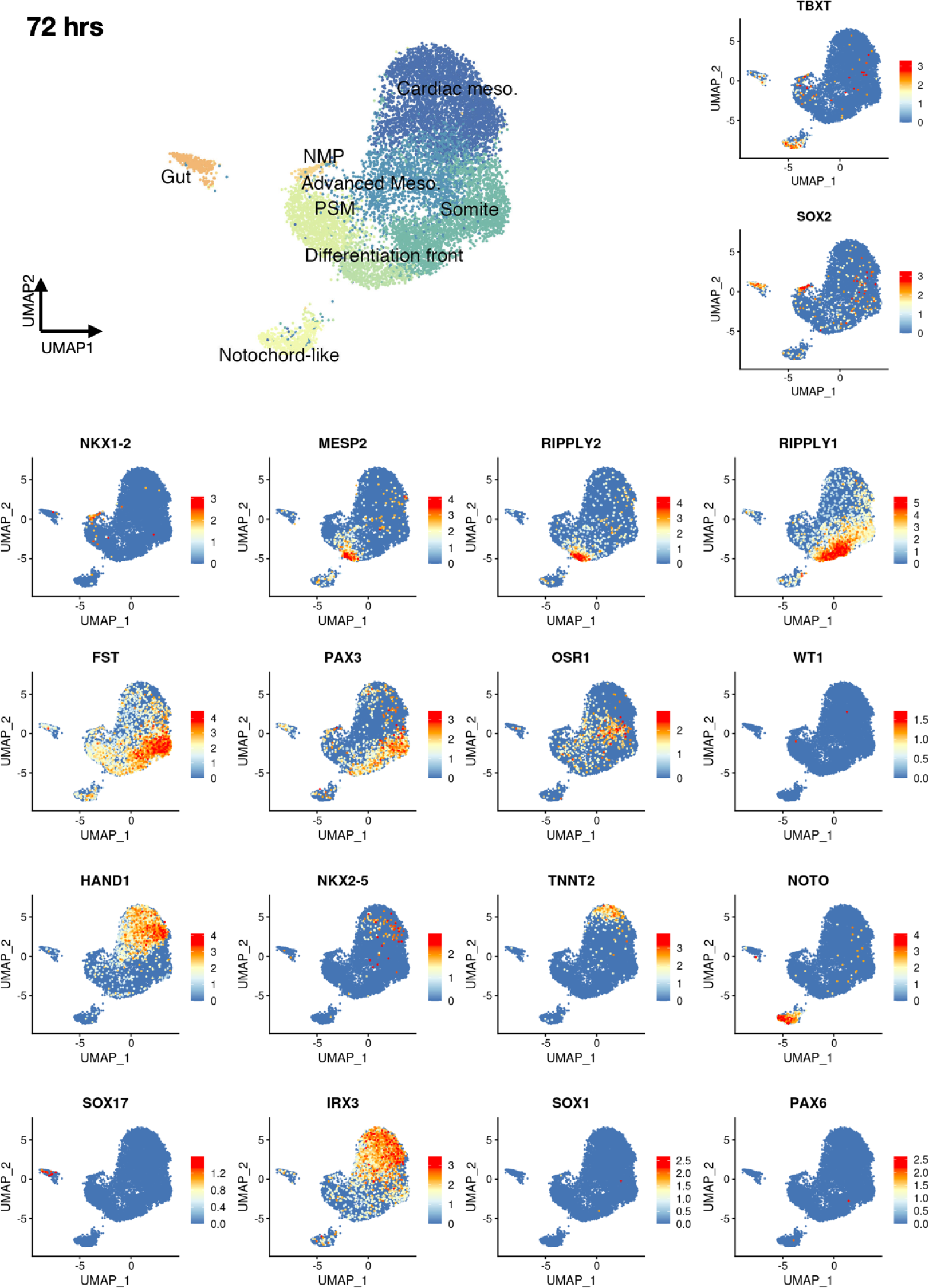
Marker gene expression and cell type annotation in human gastruloids at 72 hrs after induction. UMAP with unsupervised clustering of sc-RNA-seq profiles is shown, along with expression of marker genes used to annotate these clusters as a continuum of neuromesodermal progenitors, presomitic mesoderm, differentiation front, and somites; additionally, advanced mesoderm and cardiac mesoderm, as well as notochord-like cells and gut/endoderm-like cells. NMP, neural mesodermal progenitor; PSM, presomitic mesoderm.

**Figure S5.**
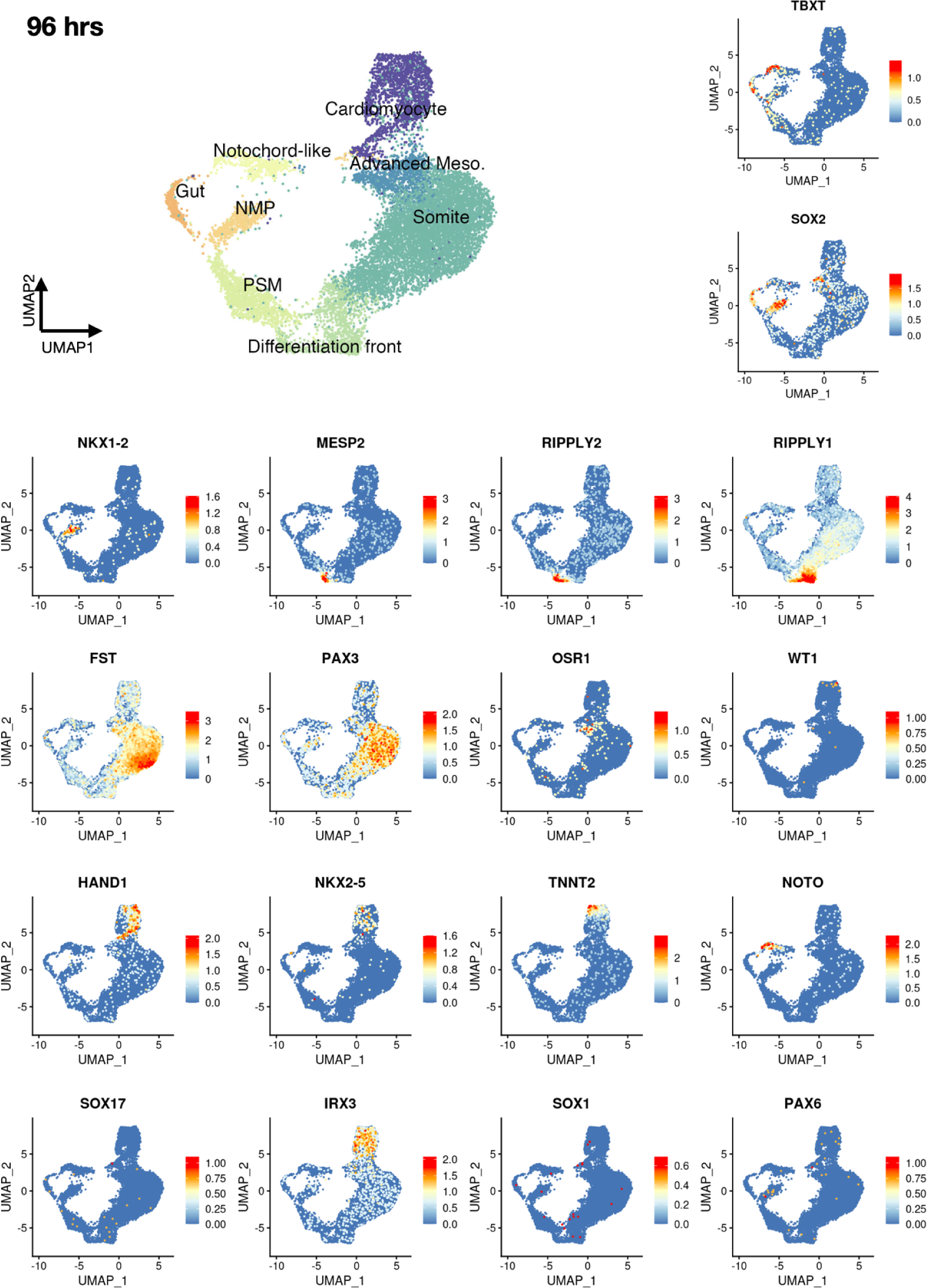
Marker gene expression and cell type annotation in human gastruloids at 96 hrs after induction. UMAP with unsupervised clustering of sc-RNA-seq profiles is shown, along with expression of marker genes used to annotate these clusters as a continuum of neuromesodermal progenitors, presomitic mesoderm, differentiation front, and somites; and additionally, advanced mesoderm and cardiac mesoderm, as well as notochord-like cells and and gut/endoderm-like cells. NMP, neural mesodermal progenitor; PSM, presomitic mesoderm.

**Figure S6.**
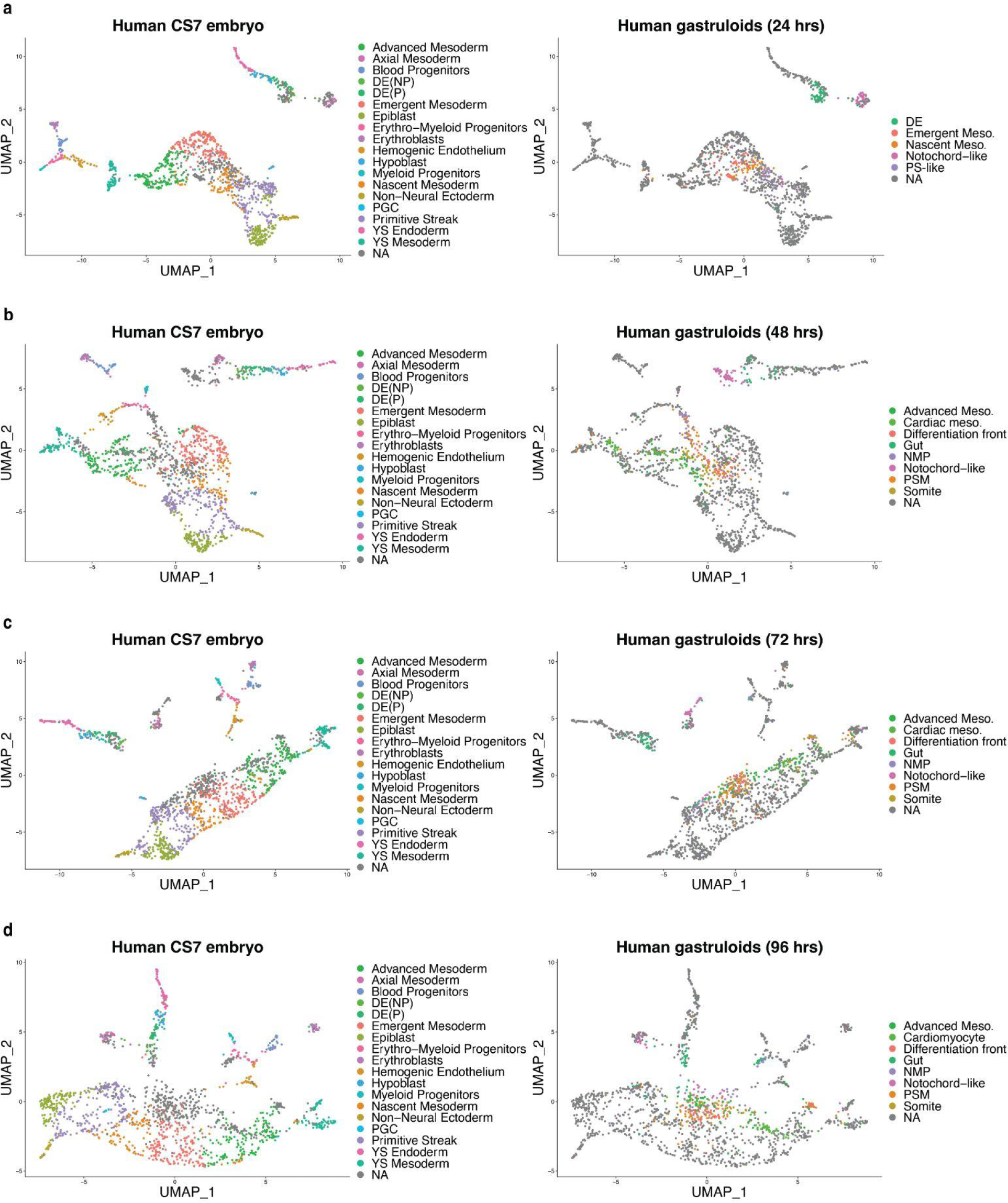
Co-embedding of human gastruloid sc-RNA-seq profiles (24-96 hrs) with published data from human embryos. Co-embedding of sc-RNA-seq profiles, labeled by annotations from human CS7 embryo^18^ (left) or human gastruloids (this study, right). NA refers to cells from the other sample in the co-embedding, which are annotated in the other column. Each row corresponds to a different timepoint of the human gastruloids from 24 to 96 hours.

**Figure S7.**
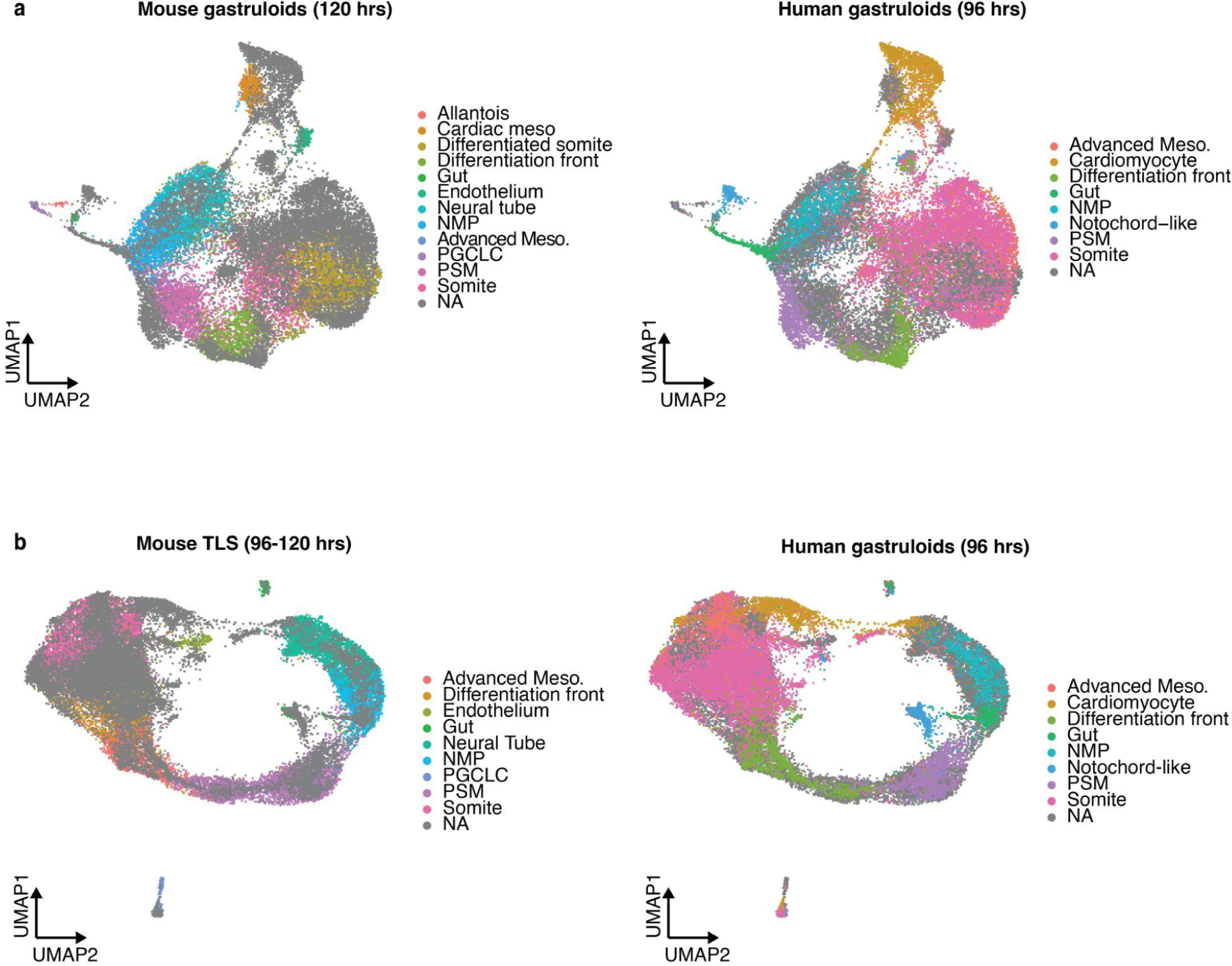
Co-embedding of human gastruloid sc-RNA-seq profiles (96 hrs) with published data from two mouse gastruloid models. **a**, Co-embedding of sc-RNA-seq profiles, labeled by annotations from 120 hr mouse gastruloids^2^ (left) or 96 hr human gastruloids (right). NA refers to cells from the other sample in the co-embedding, which are annotated in the other column. **b**, Co-embedding of sc-RNA-seq profiles, labeled by annotations from 96-120 hr mouse TLS^15^ (left) or 96 hr human gastruloids (right).

**Figure S8.**
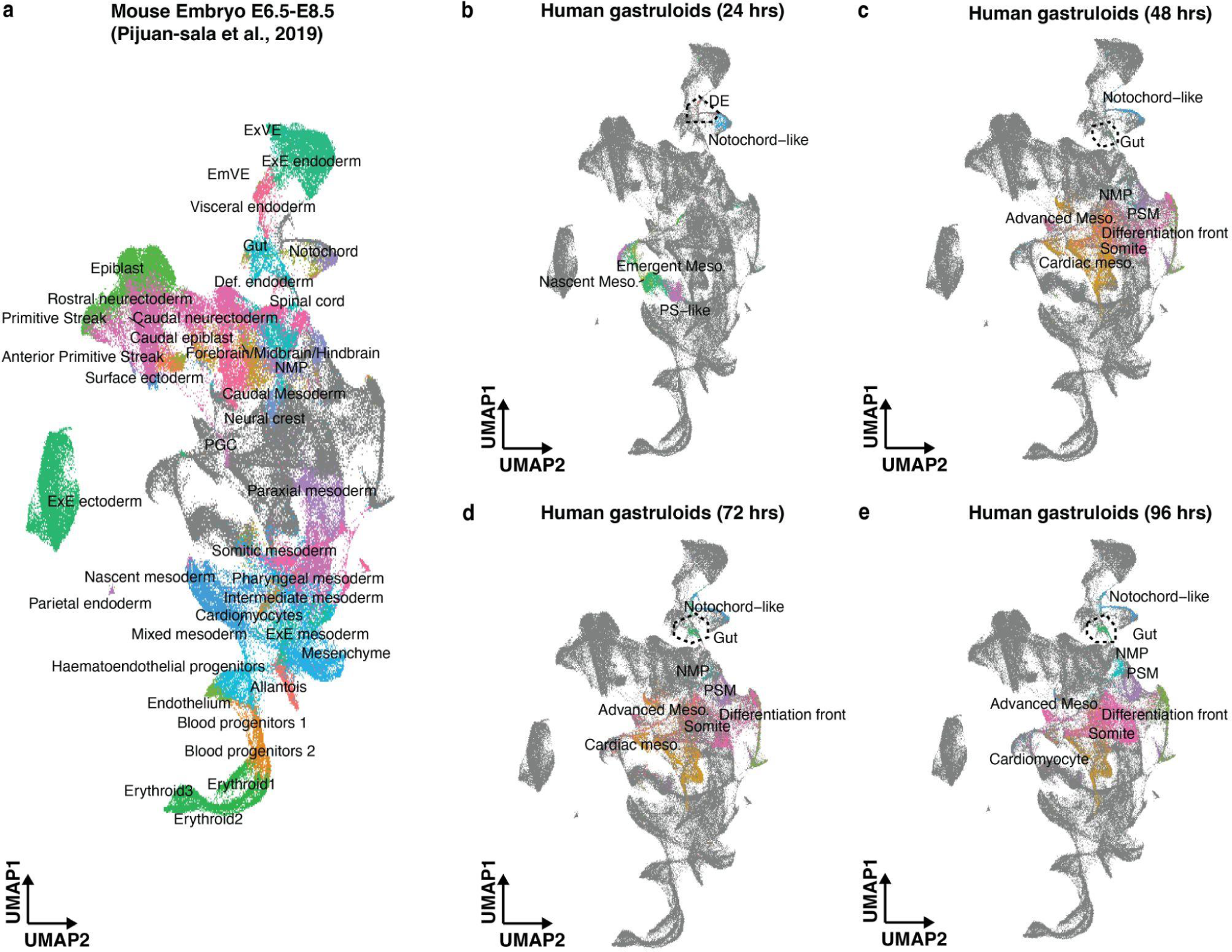
Mapping endoderm-like cells via integration of human gastruloid and mouse embryo sc-RNA-seq data. **a**, UMAP visualization of co-embedding of data from 24-96 hr human gastruloids (this study) and E6.5-E8.5 mouse embryos^51^. Mouse embryo cells are colored and annotated with labels from the original study. **b-e**, Same as panel (**a**), with human gastruloid cells from 24 hrs (**b**), 48 hrs (**c**), 72 hrs (**d**) or 96 hrs (**e**), colored and annotated with labels from this study. Cells that we have annotated as resembling definitive endoderm (DE) at 24 hrs co-localize with mouse embryonic visceral endoderm (EmVE) cells. Cells that we have annotated as gut/endoderm-like at 48-96 hrs co-localize with mouse embryo gut cells.

**Figure S9.**
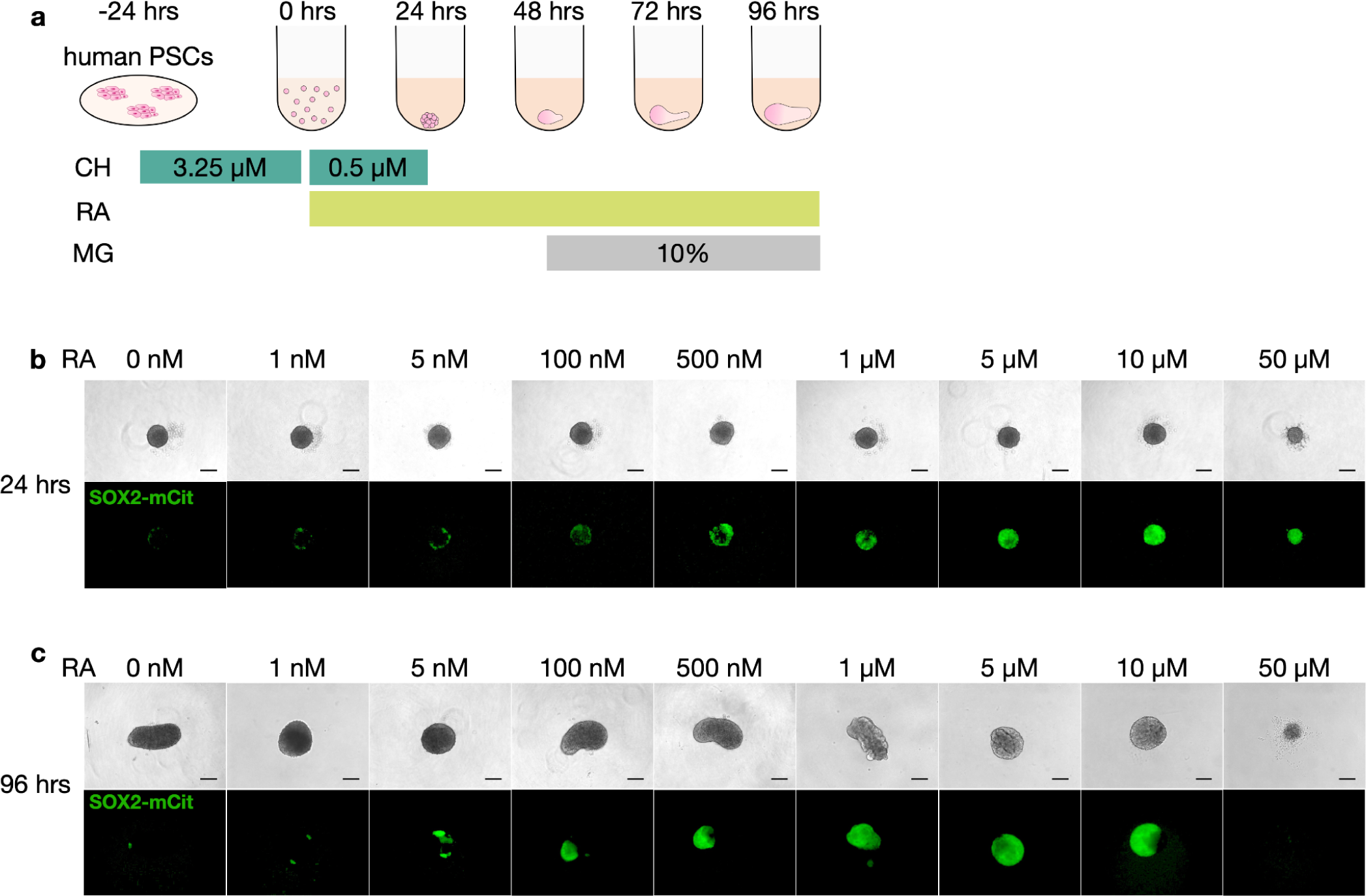
Effect of continuous introduction of retinoic acid on neural cell induction in human gastruloids. **a**, Schematic illustration of RA and Matrigel treatment on human gastruloids. RA, retinoic acid; MG, 10% Matrigel. **b**-**c**, Representative images of human gastruloids at 24 hrs (**b**) and 96 hrs (**c**). The intensity of SOX2-mCit correlated with the concentration of RA. Scale bars, 100 µm. N = 48 gastruloids for each concentration showed similar morphology and expression patterns of marker genes.

**Figure S10.**
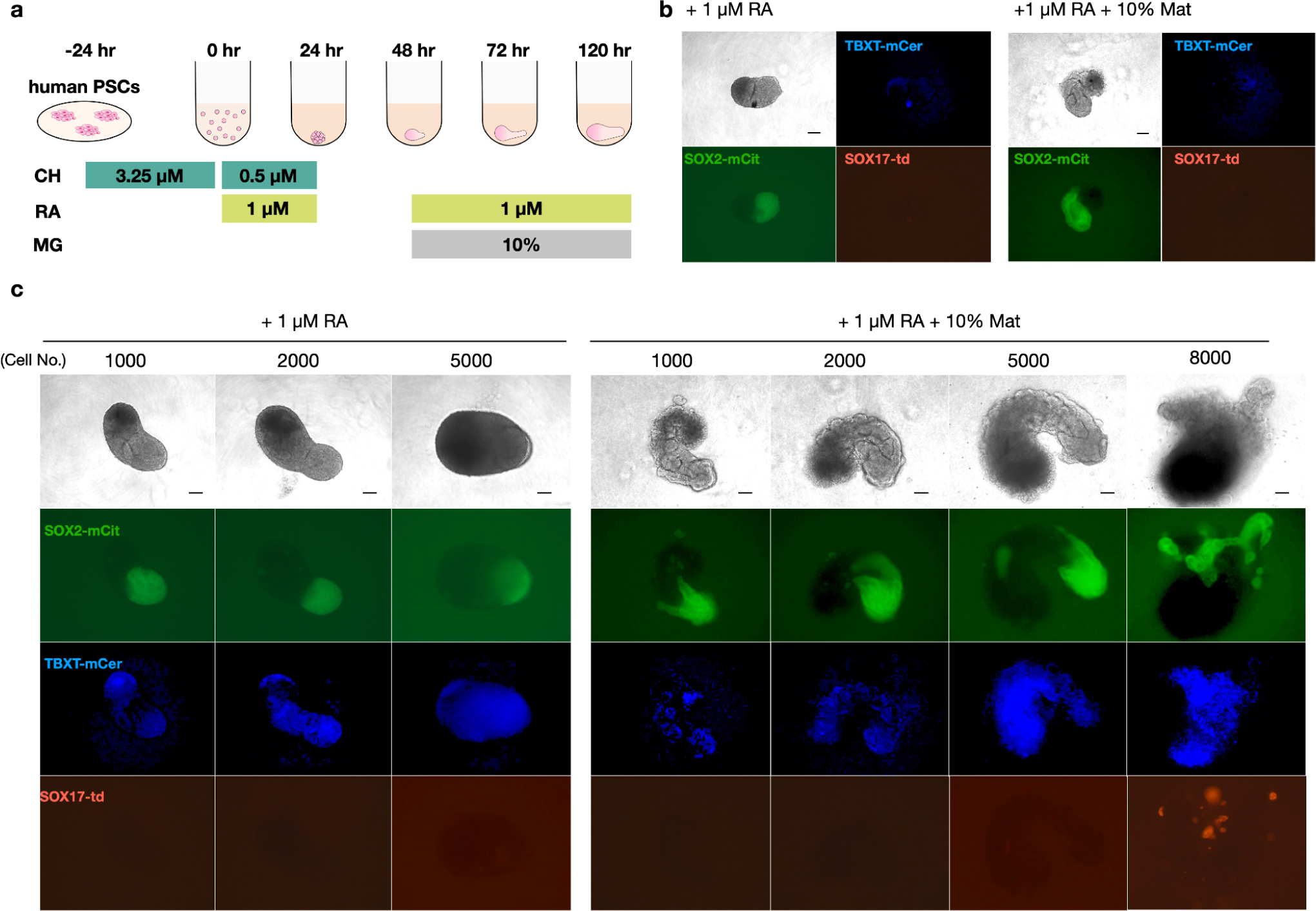
Number of seeded cells impacts human gastruloid formation, in the context of a discontinuous regimen of retinoic acid. **a,** Schematic of discontinuous regimen of RA and Matrigel treatment while inducing human gastruloids. RA, retinoic acid; MG, 10% Matrigel. **b**, Representative images of 96 hr human gastruloids induced from 400 cells. 1 µM RA (0-24 hrs and 48-96 hrs) (left) or 1 µM RA (0-24 hrs and 48-96 hrs) + 10% Matrigel (48-96 hrs) (right) were added to the medium. Scale bars, 100 µm. N = 32 gastruloids showed similar morphology and expression patterns of marker genes. **c**, Representative images of 96 hr human RA-gastruloids while varying the number of cells used for initial seeding. Scale bars, 100 µm. N = 48 RA-gastruloids showed similar morphology and expression patterns of marker genes.

**Figure S11.**
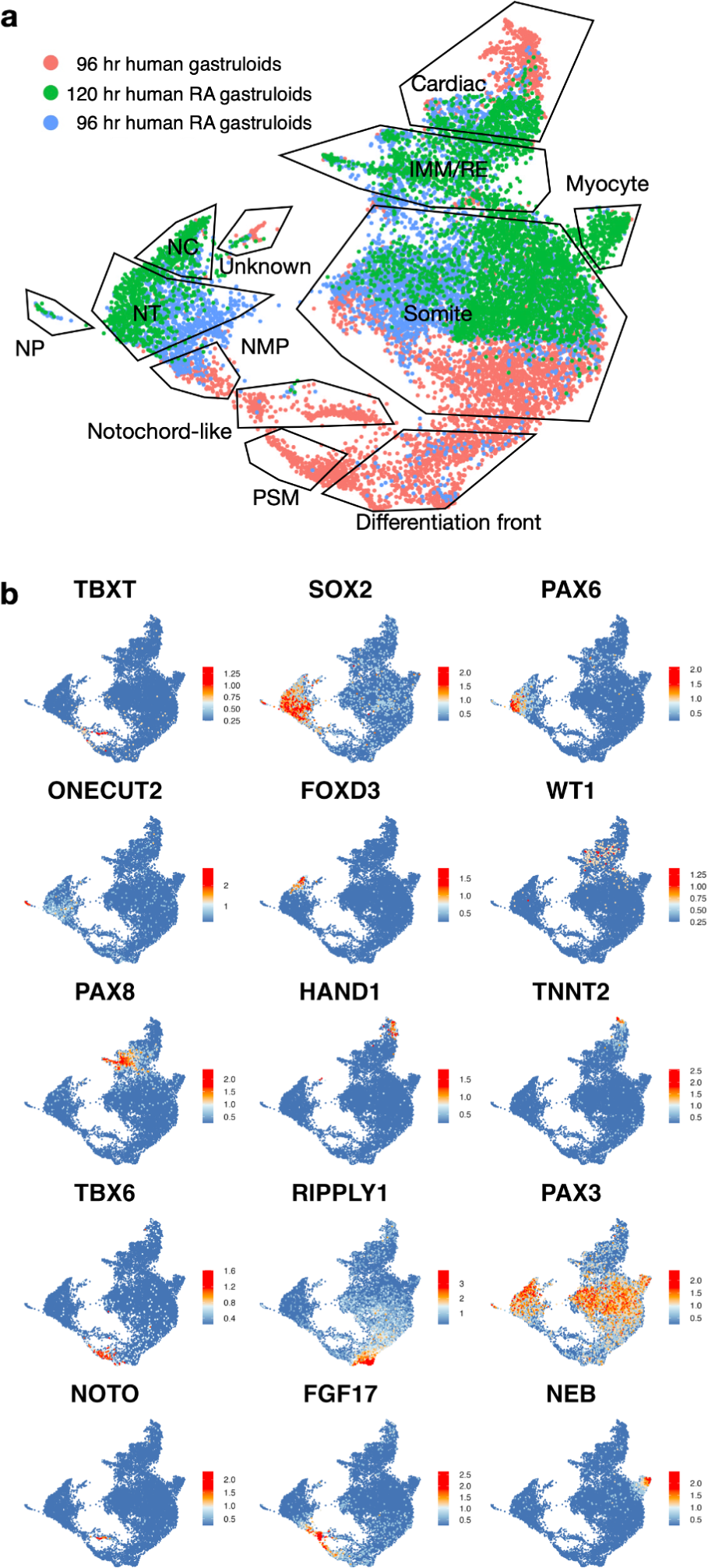
Comparison of cell types detected in human non-RA vs. RA-gastruloids. **a**, Co-embedded UMAP of sc-RNA-seq data colored by the sample of origin. Red: 96 hr non-RA-gastruloid (n = 5,000 cells). Blue: 96 hr RA-gastruloid (n = 5,000 cells). Green: 120 hr RA-gastruloid (n = 5,000). Clusters were annotated based on the expression pattern of marker genes listed in panel (**b**). NP, neural progenitors; NC, neural crest; NT, neural tube; PSM, presomitic mesoderm; IMM, intermediate mesoderm; RE, renal epithelium; NMP, neuromesodermal progenitor. **b**, Marker gene expression on co-embedded UMAP.

**Figure S12.**
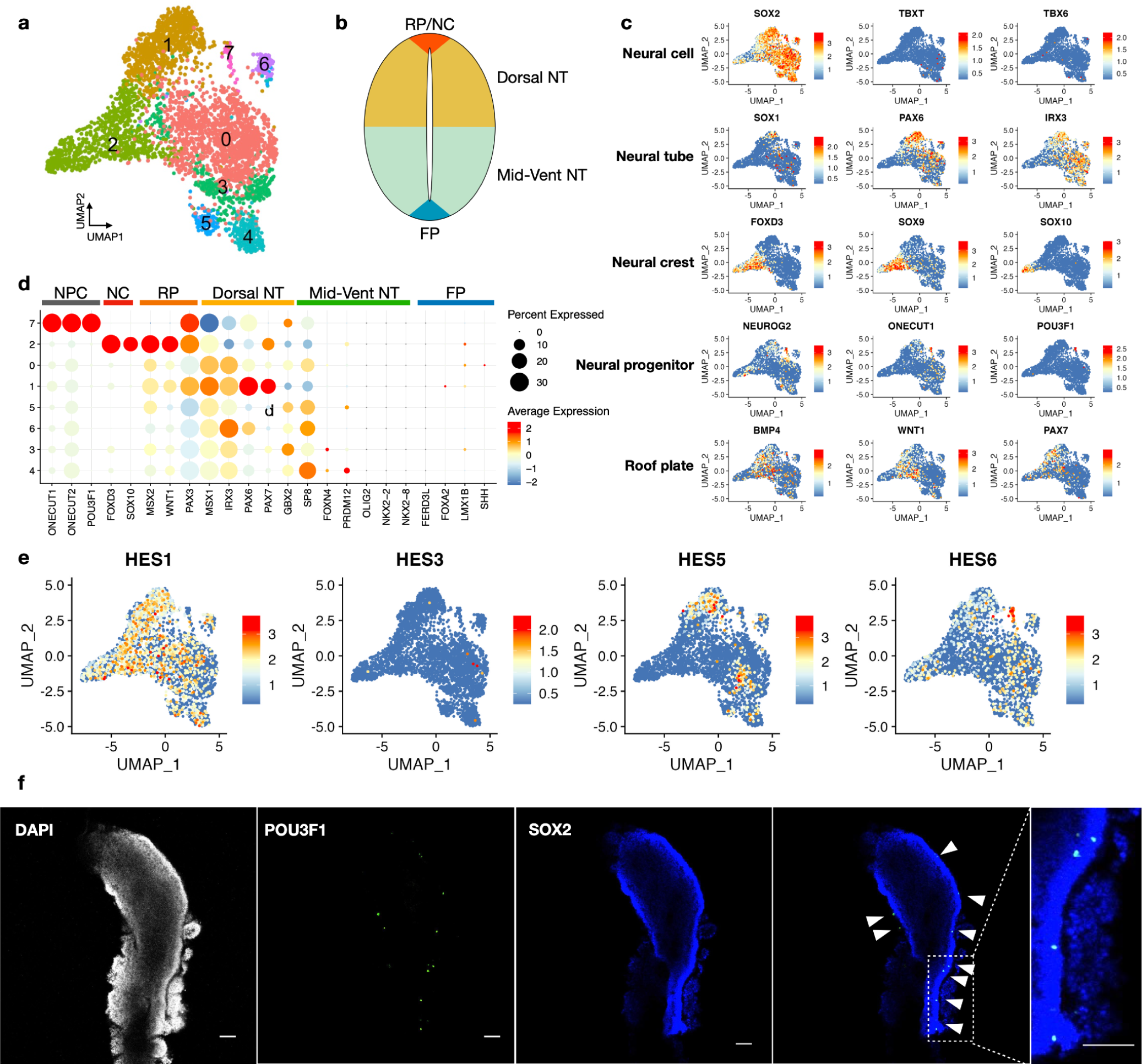
Sub-clustering and annotation of neural cells. **a**, UMAP and clustering of neural-related cells from 120 hr human RA-gastruloids. **b**, Schematic illustration of marker gene expression patterns in neural tube in relation to dorsal-ventral axis. **c**, Marker gene expression projected on UMAP shown in panel (**a**). **d**, Bubble plot of marker gene expression patterns in each cluster. **e**, Expression patterns of HES genes in neural-related cells of 120 hr human RA-gastruloid. *HES1*+/*HES*5*+* and *HES6*+ mark radial glia-like cells and neural progenitors, respectively^44^. The neural progenitors (cluster 6) are also *ONECUT1*+, *ONECUT2*+, *NEUROG2,*+, *POU3F1*+, as shown in panels (**c**) and (**d**). **f**, Immunostaining of neural progenitor cells in 120 hr RA-gastruloid. Neural progenitor cells were stained with anti-POU3F1 antibody (white) and the neural tube was stained with anti-SOX2 antibody (blue). POU3F1+ neural progenitors were sparsely distributed across the anterior-posterior axis of the neural tube-like structure. The dotted box is magnified in the right panel. Scale bar, 100 µm. N=5 human RA-gastruloids showed similar morphology and expression patterns of marker genes.

**Figure S13.**
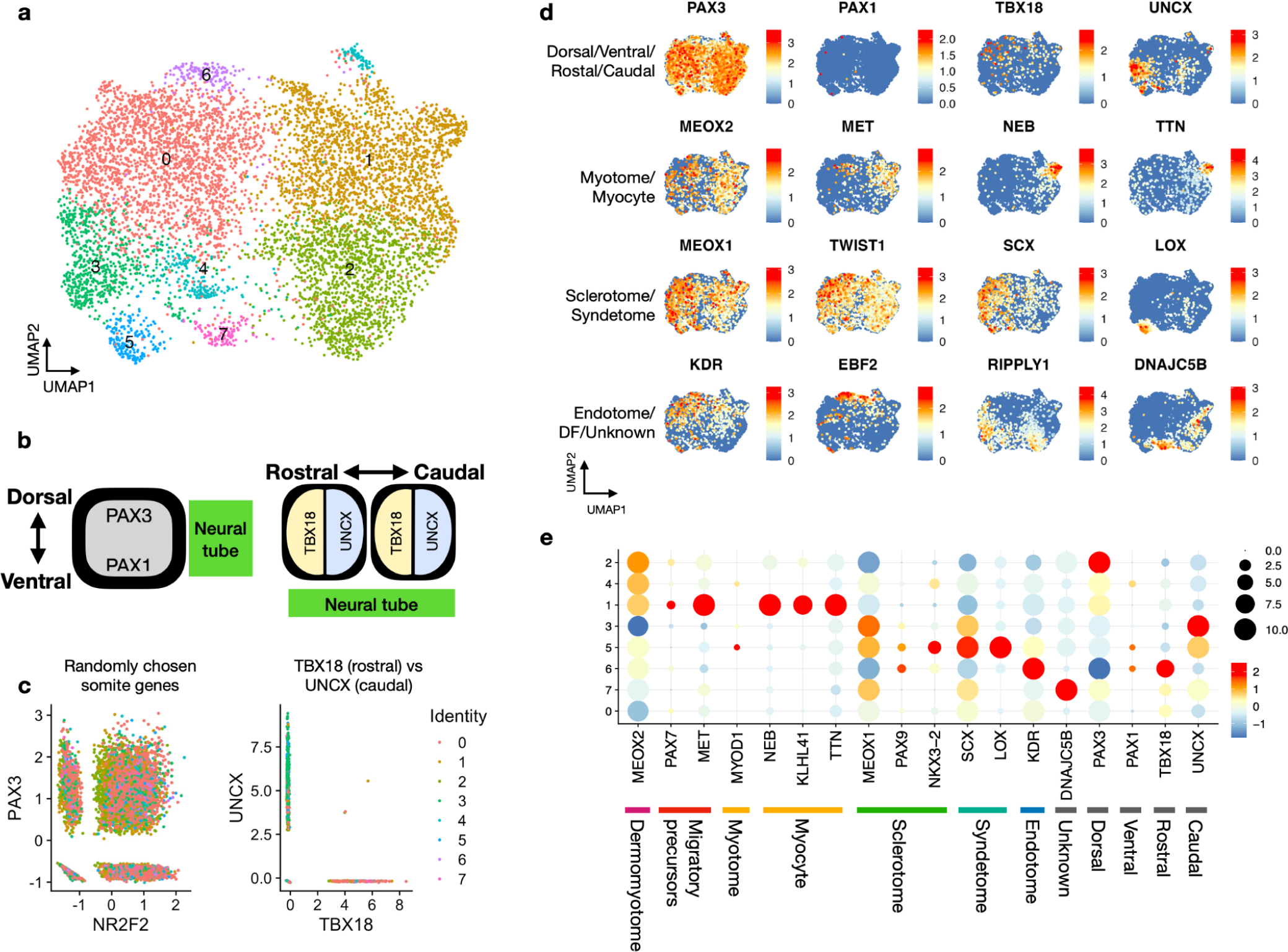
Sub-clustering and annotation of paraxial mesoderm derivatives. **a**, UMAP of sub-clustered somite, differentiation front, and myocyte cluster cells from 120 hrs RA-gastruloid sc-RNA-seq data. **b**, Schematic illustration of marker gene expression patterns in somite. **c**, Scatter plot showing scaled expression levels of randomly chosen pairs of genes (left) and *TBX18* and *UNCX* (right). The expression of *TBX18* and *UNCX*, markers of the rostral-caudal axis of somites, was mutually exclusive. **d**, Expression patterns of selected marker genes. DF, differentiation front. **e**, Bubble plot showing expression patterns of selected marker genes. DF, differentiation front.

**Figure S14.**
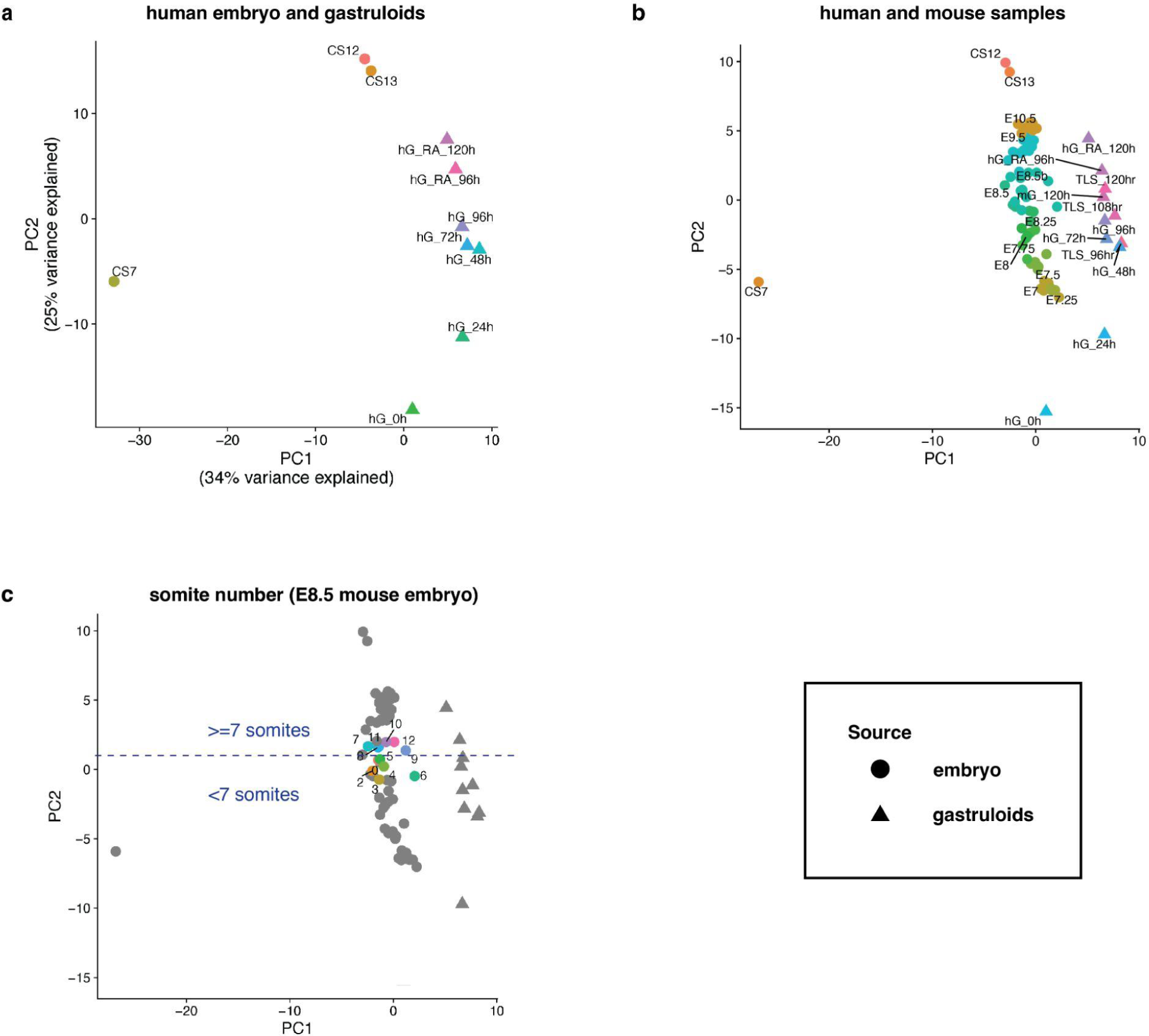
Principal components analysis (PCA) of *in vitro* models of early mammalian embryogenesis. **a**, PCA space computed on pseudo-bulk transcriptomes of pooled RA human gastruloids (96 and 120 hrs), pooled original human gastruloids (0, 24, 48, 72 and 96 hrs) and individual human embryos at CS7, CS12, and CS13. A set of 455 highly variable genes in human embryo datasets^18,47^ that also showed expression in both human and mouse gastruloid datasets was used for this analysis. PC2 is strongly correlated with developmental progression. **b**, Projection of data from pooled mouse gastruloids (120 hrs), pooled TLS (96, 108 and 120 hrs), and individual^49,50^ or pooled^51^ mouse embryos, onto the PC space defined by the analysis of human data shown in panel (**a**). For mouse embryos, E8.5 and earlier samples are pooled embryos processed by 10X Genomics sc-RNA-seq^51^, while E8.5b and later samples are individual mouse embryos processed by sci-RNA-seq3^50,52^. **c**, Same as panel (**b**), but highlighting mouse embryos from E8.5b samples that are somite-staged^50^.

**Figure S15.**
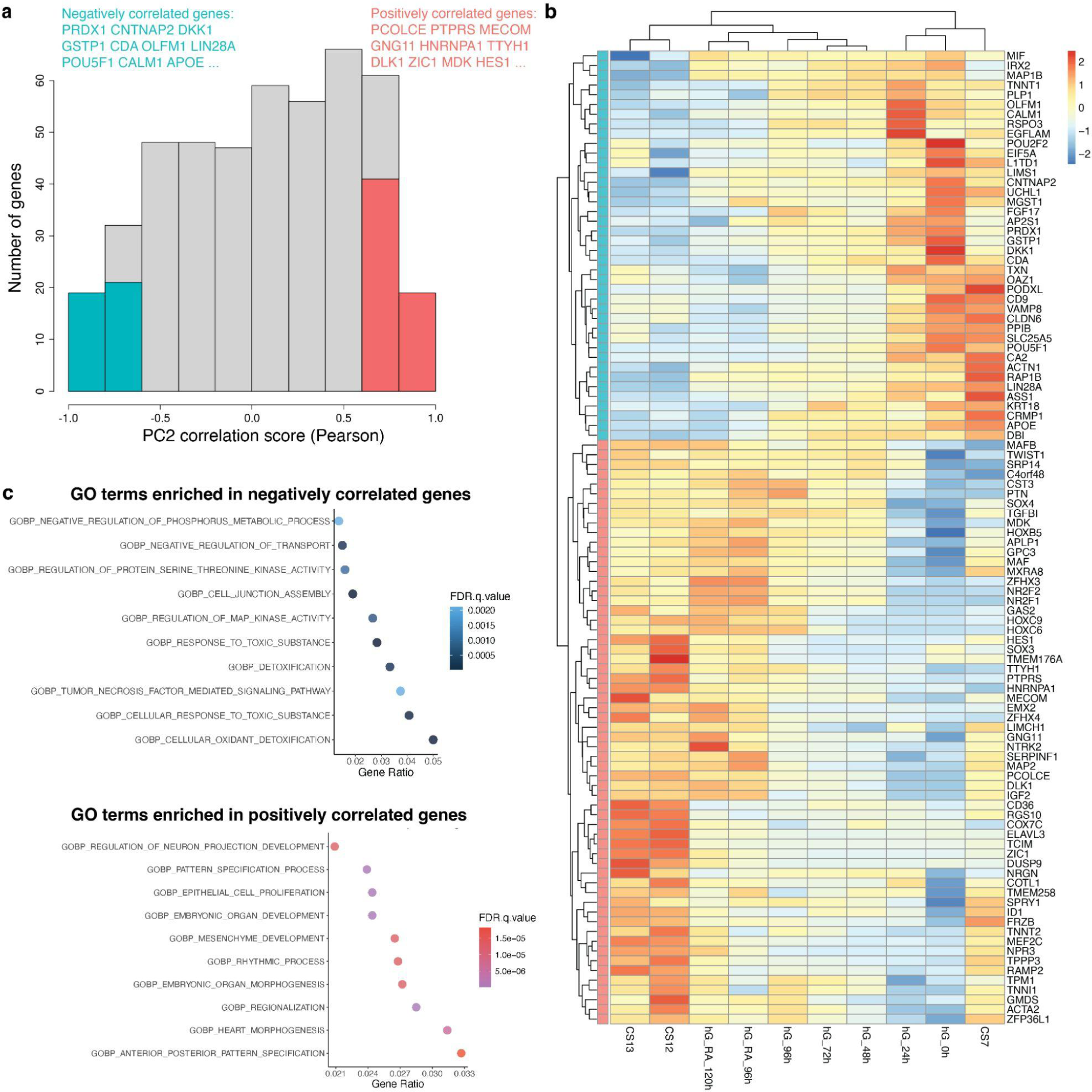
Genes that are highly correlated with developmental progression (PC2). **a**, The genes (top 100) with the highest absolute correlation scores with PC2 were selected from all 455 genes (Methods) used for PC analysis of human samples^18,47^. **b**, Heatmap shows the z-transformed expression levels of each of these top 100 genes positively (pink) or negatively (blue) correlated with PC2 in various human samples. Hierarchical clustering separated less vs. more advanced gastruloids/embryos from one another. **c**, Gene Ontology (GO) enrichment analysis (Molecular signature database v7.5.1) of the genes negatively (top) or positively (bottom) correlated with PC2^87,88^.

**Figure S16.**
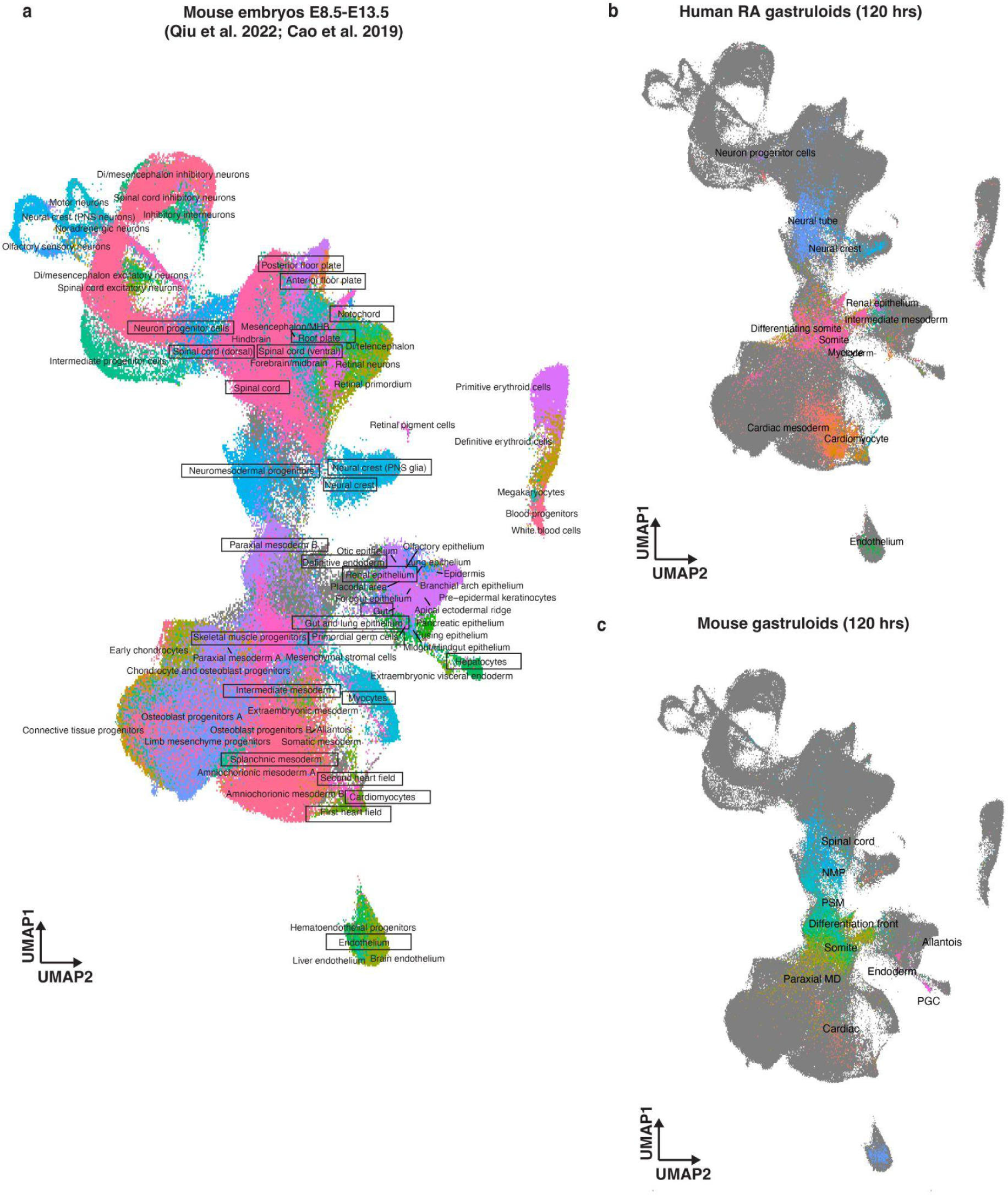
Co-embedding of single cell RNA-seq profiles from mouse embryos (E8.5-13.5), human RA-gastruloids and mouse gastruloids. **a**, Co-embedding labeled by mouse embryo cell types^49,50^. Mouse embryo cell types selected for NNLS analyses presented in **Fig. 4d** and **Fig. S17** are boxed. **b**, Co-embedding labeled by human RA-gastruloid cell types. **c**, Co-embedding labeled by mouse gastruloid^2^ cell types.

**Figure S17.**
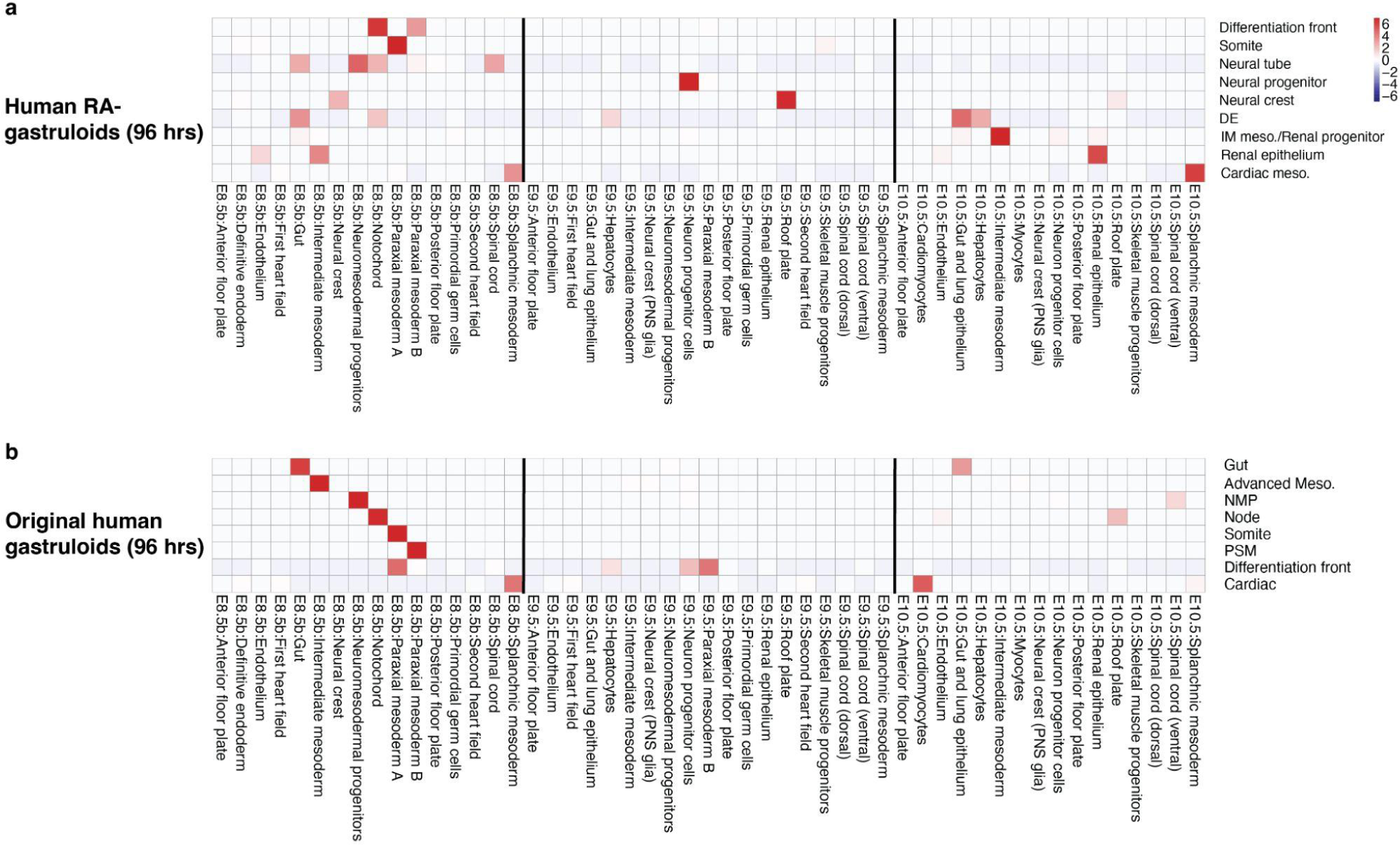
Cell type correlation analysis between human *in vitro* gastruloids vs. mouse embryos (E8.5-10.5). Correlated cell types between human gastruloids at 96 hrs and mouse embryos (E8.5-10.5)^49,50^ based on nonnegative least-squares (NNLS) regression. **a**, Heatmap of scaled correlation coefficient (z-score) between 9 cell types from human RA-gastruloids at 96 hrs vs. 16 cell types from E8.5 mouse embryos, 19 cell types from E9.5 mouse embryos and 16 cell types from E10.5 mouse embryos.^49,50^ **b**, Same, as panel (**a**), but comparing 8 cell types from the original human gastruloids^1^ at 96 hrs to the same sets of mouse cell types. At 96 hours, the human RA-gastruloids appear more advanced but also more desynchronized than the original human gastruloids.

**Figure S18.**
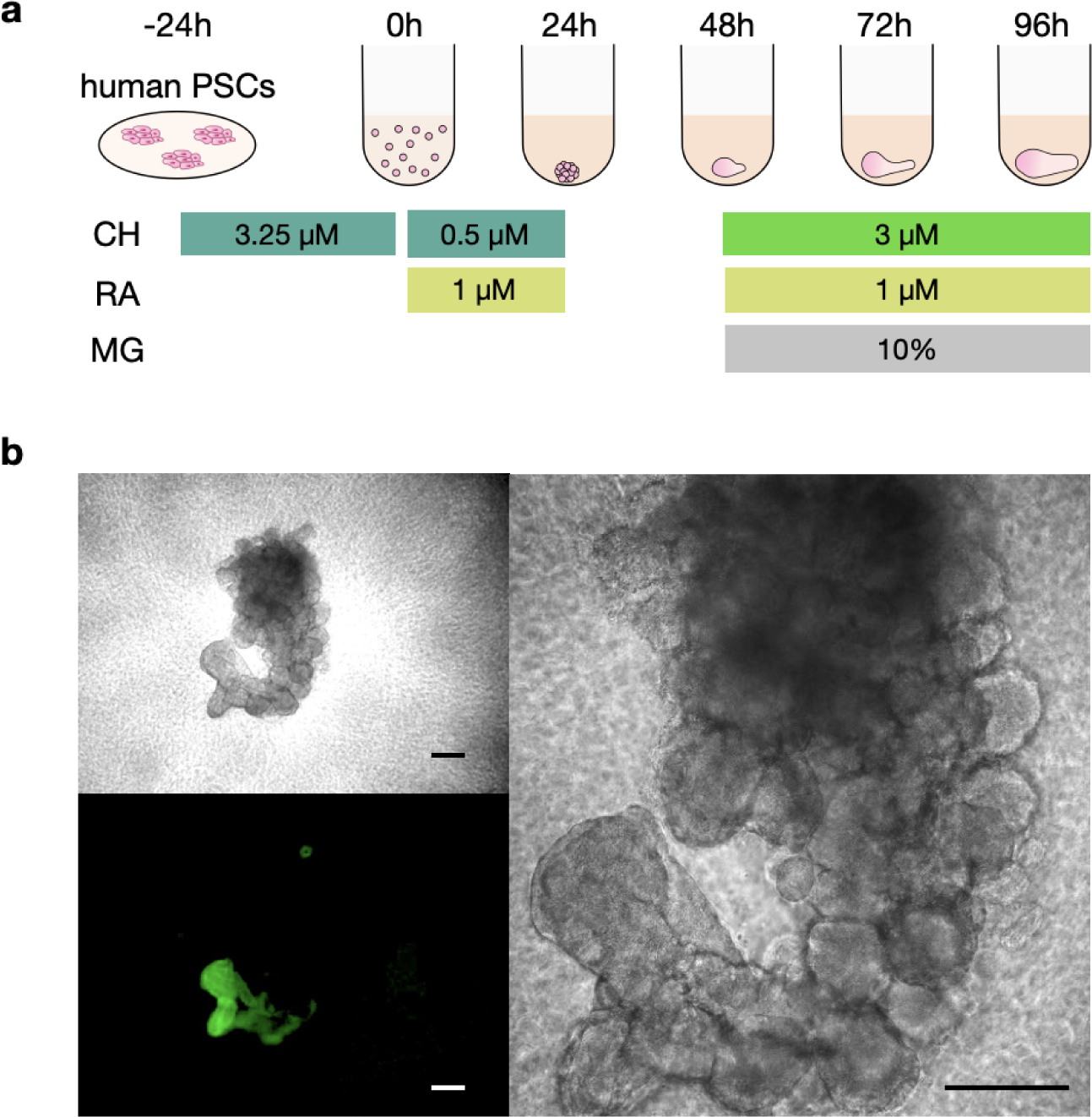
Hyper-activation of WNT signaling in human RA-gastruloids. **a**, Schematic of Hyper-activation of WNT signaling in RA-gastruloid. This protocol was modified to add CHIR back into the system at 48 hrs. CHIR, CHIR99021; MG, Matrigel; RA, Retinoic acid. **b**, Representative morphology of CHIR-treated 120 hr RA-gastruloid. Bar, 100 µm. N=192 human RA-gastruloids generated under the modified protocol showed similar morphology and marker gene expression.

**Fig. S19.**
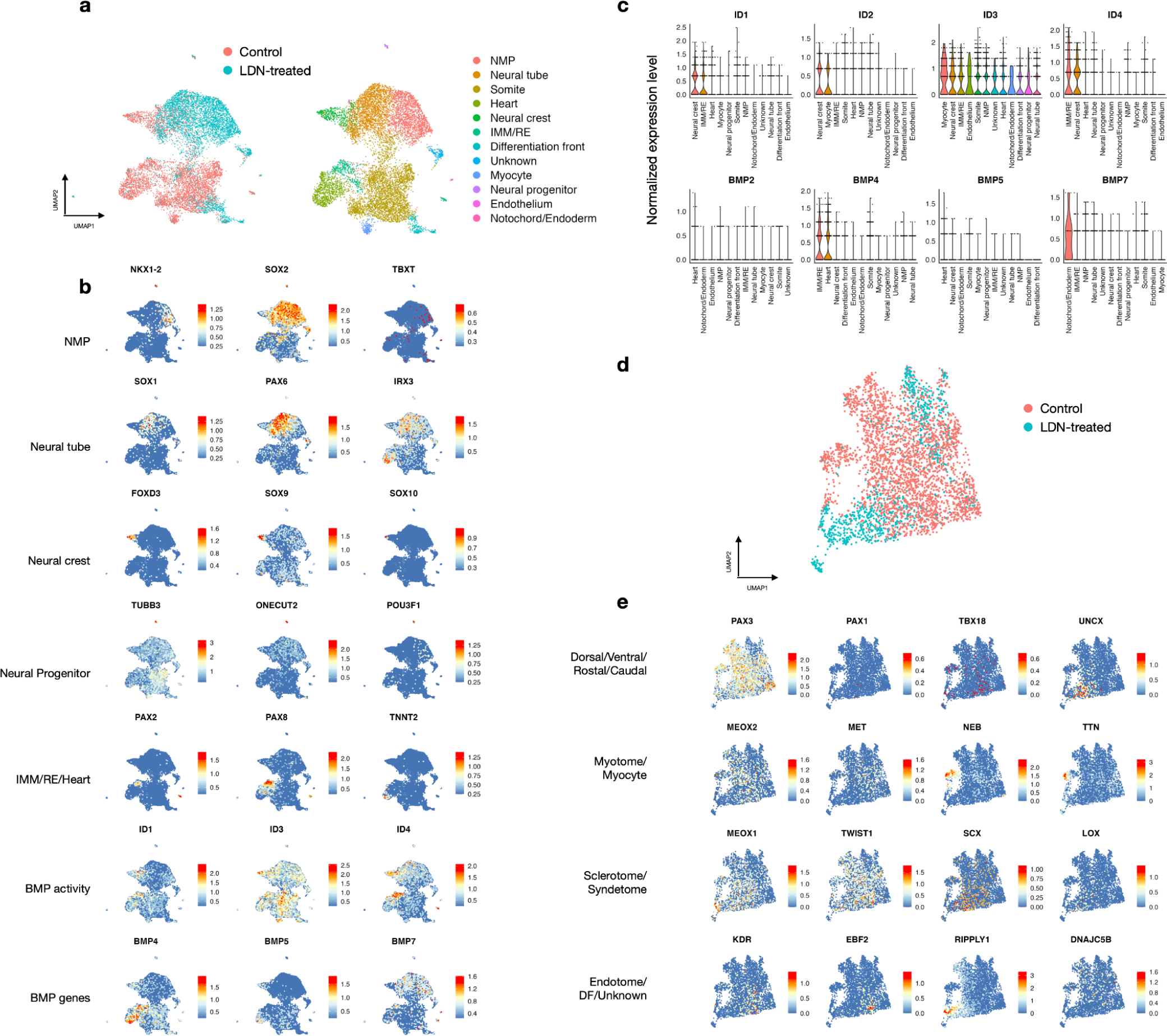
Lineage marker and BMP-related gene expression patterns in LDN-treated RA-gastruloids. **a**, Co-embedded UMAP of sci-RNA-seq data from LDN-treated RA-gastruloids vs. untreated RA-gastruloids, labeled by source (left) or cluster annotation (right). **b**, Marker gene expression patterns in co-embedded UMAP. **c**, Violin plots showing the ID and BMP gene expression patterns for each cell type. Cell types along x-axes are sorted by the mean expression levels of the gene (left, high; right, low). **d**, Subcluster analysis of somite cluster shown in panel (**a**). **e**, Marker gene expression patterns in co-embedded UMAP of somite cluster cells.

**Figure S20.**
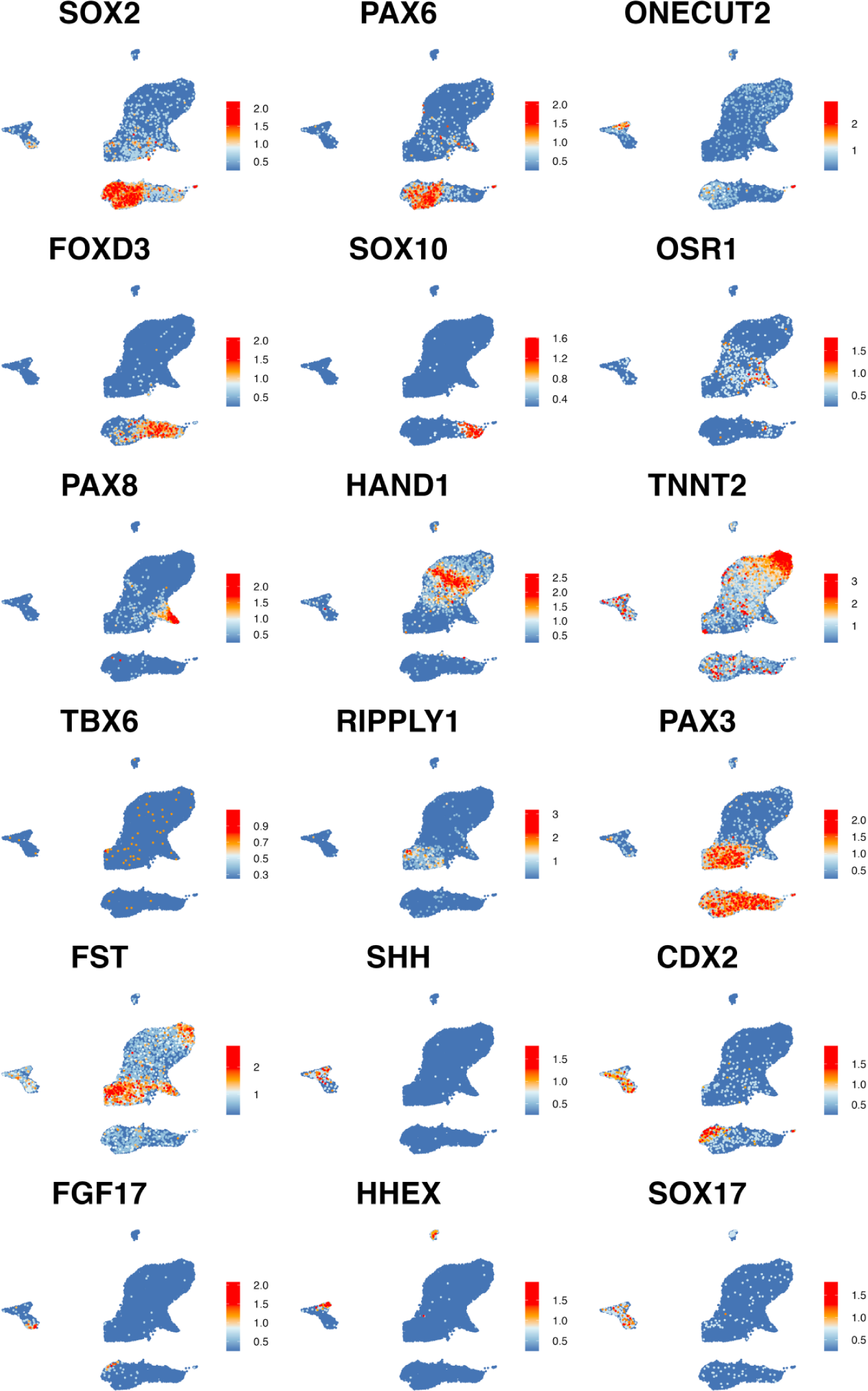
Marker gene expression in NTC and genetically-perturbed human RA-gastruloids. Marker gene expression in NTC and genetically-perturbed human RA-gastruloid on the co-embedded UMAP shown in Fig. 6d. SOX2, neural cells; PAX6, neural tube; ONECUT2, neural progenitor; FOXD3 and SOX10, neural crest; OSR1, intermediate mesoderm; PAX8, renal epithelium; HAND1, cardiac mesoderm; TNNT2, cardiac myocyte; TBX6, presomitic mesoderm; RIPPLY1; differentiation front; PAX3 and FST, somite; FGF17, definitive endoderm; HHEX, gut; SOX17, pan-endoderm.

**Figure S21.**
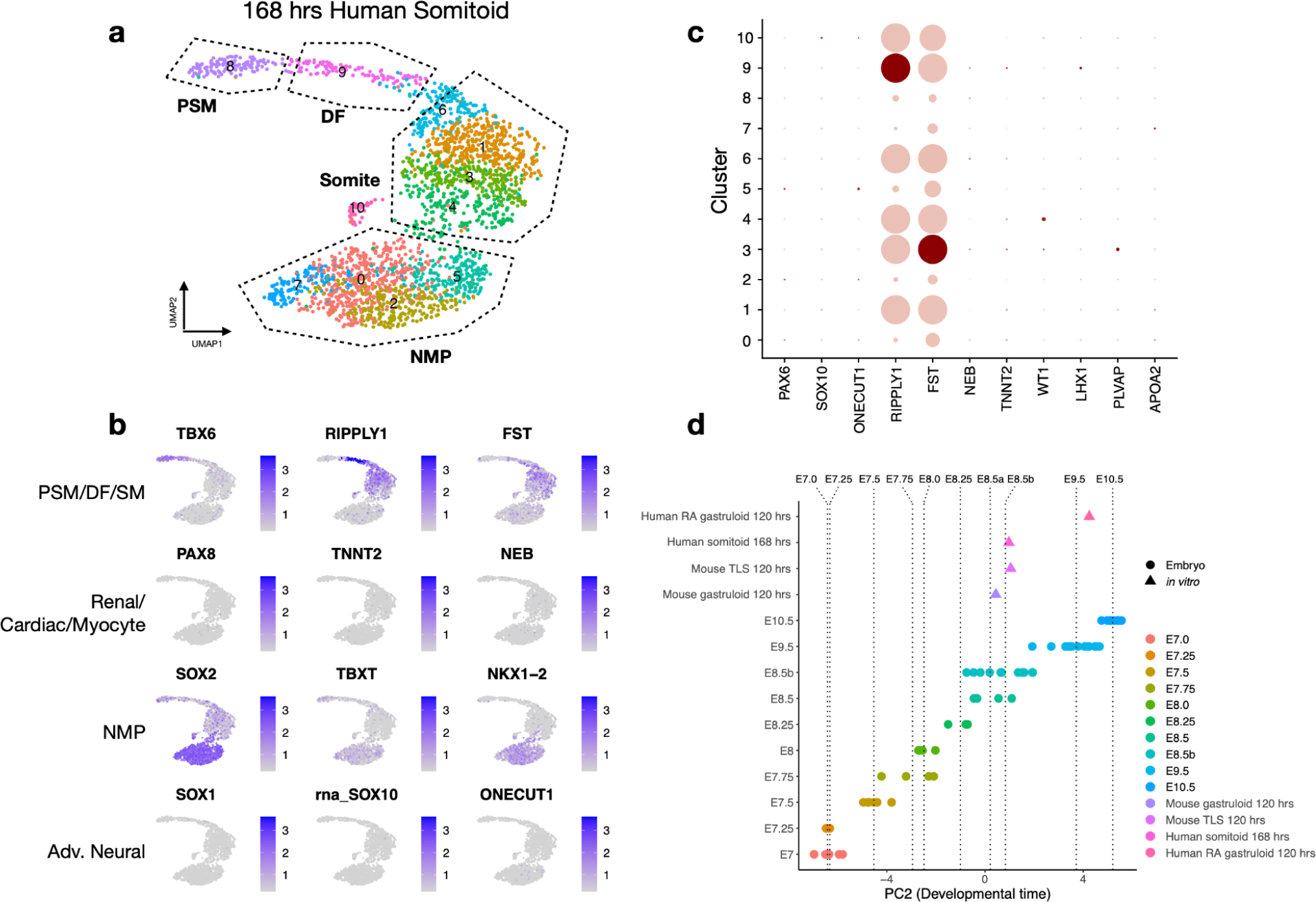
Transcriptomic evaluation of human somitoids. **a**, UMAP clustering of sc-RNA-seq data from 168 hr human somitoids^16^. **b**, Marker gene expression in UMAP of 168 hr human somitoids^16^. **c**, Dot plot of marker gene expression in somitoids. Markers are for advanced cell types observed in 120 hr human RA-gastruloids. **d**, Cross-species staging of pseudo-bulk transcriptomes of pooled human 168 hrs somitoids with the samples shown in Fig. 4c, projected on the second principal component of the PCA analysis shown in **Fig. S14a**.

